# Deciphering the zebrafish hepatic duct heterogeneity and cell plasticity using lineage tracing and single-cell transcriptomics

**DOI:** 10.1101/2025.01.09.631719

**Authors:** Jiarui Mi, Lipeng Ren, Ka-Cheuk Liu, Lorenzo Buttò, Daniel Colquhoun, Olov Andersson

## Abstract

Despite the liver’s recognized regenerative potential, the role of the hepatic ductal cells (a.k.a. biliary epithelial cells), its heterogeneity, and functionality remain incompletely understood in this process. This study provides a comprehensive examination of the molecular and cellular mechanisms underpinning liver ductal development and liver regeneration in zebrafish, with a spotlight on the functional roles of *her* family genes in these processes. Using state-of-the-art knock-in zebrafish models and single-cell transcriptomics we reveal the differential expression patterns of the different *her* genes, of which *her2*, *her6*, and *her9*, were identified as specific molecular signatures for distinguishing different ductal cell types with unique morphology and spatial distribution. Particularly, *her9* serves as a pan-ductal marker and shows responsiveness to the synergistic effect of Notch and BMP signaling. By analyzing multiple single-cell RNA-seq datasets, we identify numerous ductal markers which are functional proteins for ductal integrity, and most notably CRISPR mutagenesis demonstrates that *her9* is essential for hepatocyte recovery. Using multiple transgenic and knock-in zebrafish lines and genetic fate mapping, we provide a detailed characterization of the ductal remodeling process under development and extreme loss of intra-hepatic duct, highlighting the remarkable ductal cell plasticity. Single-cell transcriptomics of lineage-traced *her9*-expressing liver ducts in static and regenerative states uncover distinct cell clusters with unique molecular signatures and morphology, reflecting the liver’s regenerative dynamics and highlight relevant key biological processes that could be leveraged to expedite liver regeneration.

## Introduction

The liver stands out as an organ with exceptional regenerative capacity, capable of restoring its function and mass following most injuries. In instances of partial hepatectomy or liver injury, the liver primarily relies on the proliferation of residual hepatocytes to restore its mass and function^1^. However, in cases of extreme hepatocyte loss, such as those encountered in chronic or end-stage liver diseases, the liver’s regenerative mechanisms are severely challenged^2^. Therefore, this has prompted researchers to explore alternative sources of liver regeneration.

Recent studies in mouse and zebrafish revealed that the liver’s regenerative capacity is multifaceted, involving not only the proliferation of existing hepatocytes but also the transdifferentiation of biliary epithelial cells (BECs) into hepatocytes under certain conditions^2–5^. These indicate the markedly inherent cell plasticity of BECs, which allows them to adapt and respond to various stimuli and environment cues^6–8^. In early zebrafish liver organogenesis, the hepatocytes and BECs are fully differentiated from the progenitor, the hepatoblast, by around 50 hours post-fertilization (hpf)^9–12^. Interestingly, several studies showed that the BECs serve as a reservoir, which can de-differentiate into a bipotent hepatoblast-like state and can further re-differentiate into hepatocytes and BECs upon liver repair in zebrafish^13–15^. Although several key signaling pathways (such as PI3K-AKT-mTOR and Notch) and regulatory mechanisms and molecules (such as *sox9b*), have been extensively studied^14, 16^, several questions remain unanswered. For example, the BECs involved in transdifferentiation are intra-hepatic ductal cells labelled by Notch-responsive *tp1* element, however, such cells are organized in a complex ductal network with no luminal structure throughout the zebrafish lifespan^17, 18^. The mature luminal ducts are poorly defined although pivotal studies indicated potential ductal heterogeneity in the zebrafish liver^19, 20^.

In the neighboring organ, the pancreas, we recently identified multiple cells of origin for the ductal system. Such complex ductal development pattern is reconciling with the formation of other pancreatic cell types to maintain the organ integrity and whole-body homeostasis^21, 22^. Inspired by our prior observations, here, we strive to answer the aforementioned fundamental questions regarding the liver duct heterogeneity as well as the developmental dynamics of the hepatic ductal trees using cutting-edge knock-in zebrafish lines and lineage tracing^23^. We also performed single-cell transcriptomics of cells within the pan-ductal lineages for a comprehensive understanding of the hepatic ductal response during liver regeneration.

## Results

### *her9* serves as a pan-ductal marker in zebrafish liver throughout the lifespan

We first examined a publicly available single-cell RNA-sequencing (scRNA-seq) dataset from adult zebrafish livers, isolated without enrichment for specific cell types, as reported by Morrison *et al*.^19^. This unbiased approach is essential for uncovering potential ductal cell types that have not yet been characterized (Supplementary Fig. 1A -C). After performing standard dimensionality reduction and cell clustering, we discerned three distinct clusters of cholangiocytes with elevated expression of typical ductal markers (*anxa4*, *agr2*, and *tm4sf4*). Cluster 1 was notable for its expression of the *hes5* family genes, including *her2* and *her15.1*. Cluster 3 displayed selective expression of *krt4*.*1* (which is annotated as *krt4* in the current reference genome GRCz11/danRer11). Cluster 2 exhibited co-expression of ductal genes and genes typical for hepatocytes (*tfa* and *cp*), which could indicate either contamination from doublet cells containing hepatocytes or the presence of transitional cells at the boundary between ductal cells and hepatocytes. Importantly, the differential expression patterns suggest that hepatic ductal cells can be categorized into more than one subpopulation.

Our previous study identified *vasnb*, a gene encoding a membrane protein, as a universal marker for ductal cells in the zebrafish pancreas^21, 23^. Immunofluorescent staining using the anti-Vasnb antibody provided clear visualization of the cell membrane throughout the entire hepatic ductal tree^24^. This approach allowed us to gain a comprehensive view of the ductal structures and provided insights into the cellular composition of the hepatic ducts. To elucidate the spatial distribution of different *her* genes, we first employed our 3’ knock-in technique to create *her9* knock-in zebrafish line^23^. We integrated a p2A-EGFP-t2A-CreERT2 genetic cassette just upstream of the *her9* gene’s STOP codon (Fig. 1A). This cassette is transcribed in tandem with the *her9* gene and subsequently self-cleaves into EGFP and CreERT2, enabling both cell labeling and lineage tracing. In 6-day post-fertilization (dpf) fish larvae, genetically fluorescent labeling was observed in both the extra-hepatic ducts (EHD) and intra-hepatic ducts (IHD), while the gallbladder was absent of fluorescence. Fluorescent imaging revealed that the *her9* knock-in signals are distributed in the whole biliary ductal tree in 6 dpf larvae (Fig. 1B and B’). This expression pattern persists as the fish mature into juveniles and adults, as depicted in Fig. 1C – E, Supplementary Fig. 4.

**Fig. 1.**
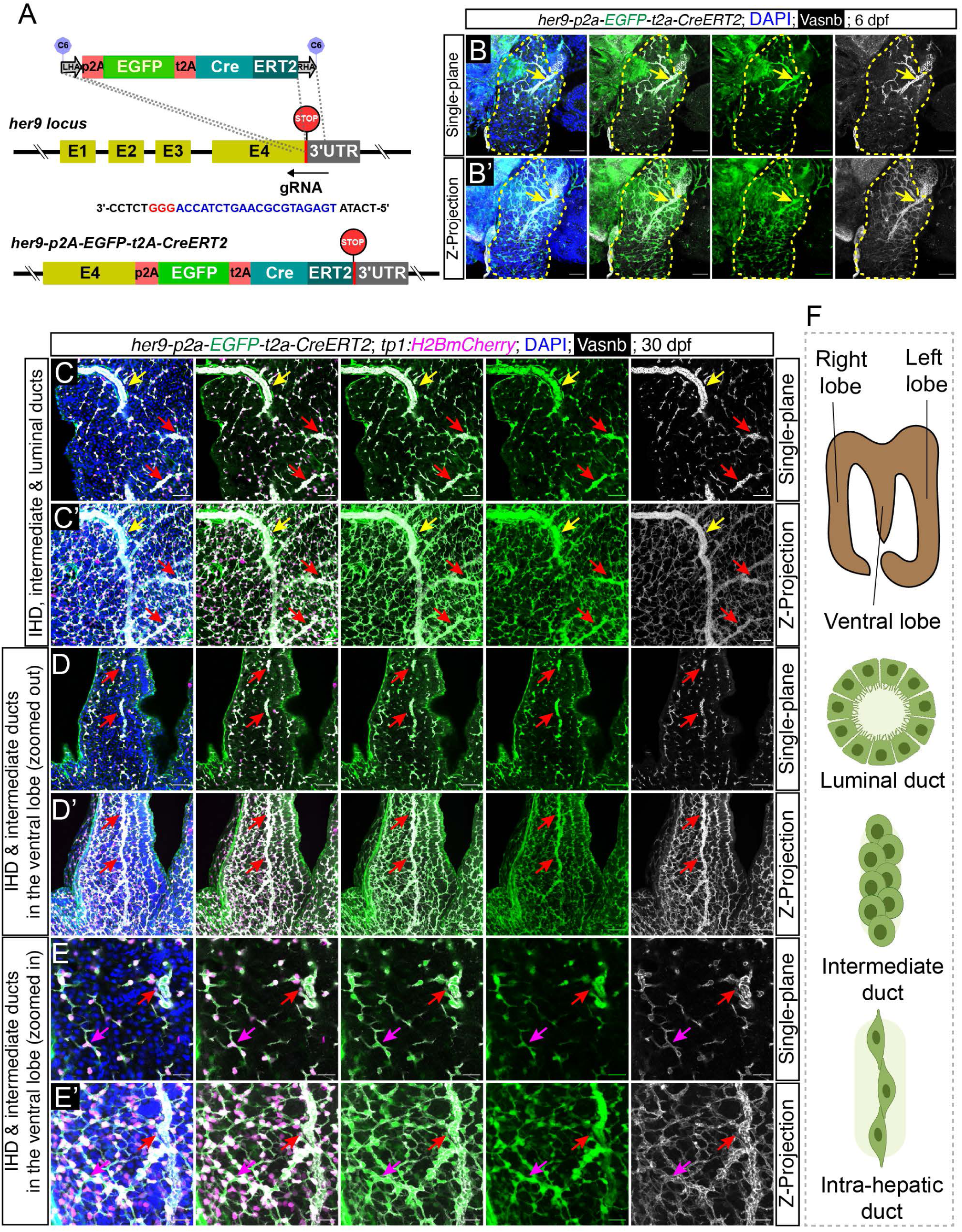
The design and characterization of a knock-in line at the *her9* locus. (A) The design of donor dsDNA templates and gRNA sequence for the construction of a knock-in zebrafish line at the *her9* locus. The nucleotide sequence in blue indicates gRNA, whereas the nucleotide sequence in red indicates the PAM sequence. (B) Representative single-plane (B) and Z-projection (B’) <u>confocal images</u> showing the *her9* labeling in hepatic biliary ductal tree, stained by a pan-ductal antibody against Vasnb, in the larval zebrafish liver. The yellow dashed line indicates the liver, and the yellow arrow indicates the large luminal duct that forms the stem of the ductal tree. (C-E) Representative single-plane (C,D,E) or Z-projection (C’,D’,E’) confocal images for the visualization of *her9* expression pattern in different hepatic regions using juvenile *TgKI(her9-p2A-EGFP-t2A-CreERT2)* homozygous knock-in reporter line. (F) The schematics show the liver lobe structure and three types of biliary ductal cells with distinct morphological structure and spatial distribution. The arrows indicate the ducts with different morphology and spatial distribution, i.e., luminal duct (yellow), intermediate duct (red), and IHD (magenta). Scale bars, 100 μm (B-D) and 40 μm (E).

We also visualized the ductal trees in isolated liver from *her9* knock-in juvenile fish. Through systematic scanning of the whole liver, we aimed to identify *her9*^+^ cells with diverse morphologies and positions. In proximity to the hilum, we observed *her9*^+^ luminal ducts accompanied by cellular sheets that extend towards the liver parenchyma, where the peripheral fibroblast-looking ductal cells are marked by the Notch-responsive *Tg(tp1:H2BmCherry)* (Fig. 1C and C’). The *tp1* element, derived from the Epstein-Barr virus and featuring consecutive RBPJ binding sites, has been extensively utilized to label Notch-responsive cells across various organs in zebrafish^17, 25^. Notably, as the luminal ducts progress, they become increasingly slender, transitioning into clusters of cells that eventually merge with the IHD. Within these clustered regions, the ductal cells exhibited a round shape, lacking distinct apical-basal polarity. This pattern was consistently observed, and especially obvious in the ventral lobe, where the main luminal duct is centrally located with the radiating branches that extend outwards, connecting to the IHD (Fig. 1D-E). Consequently, we classified this area, characterized by cell clusters, as the intermediate duct, reflecting its transitional cellular morphology and ductal structure (Fig. 1F). The *her9* expression pattern was found to be consistent in older fish (juvenile and adults), with frequent formation of intermediate ducts in both the left and right lobes, as depicted in Supplementary Fig. 2 and 4A -D.

### Differential spatial expression pattern of *her2* and *her6* in the hepatic duct

The IHD cells are widely recognized as Notch-responsive ductal cells due to their sensitivity to the inhibition of Notch signaling. Significantly, the *Tg(tp1:H2BmCherry)* signal was robust in the IHDs but reduced as the ducts expanded (in the intermediate duct) and was greatly diminished in the major ducts. This indicates a gradient of Notch signaling intensity, decreasing from the peripheral to the central ductal regions within the liver.

Considering the intricacies of Notch signaling and the multiplicity of *hes* family genes, the *tp1* transgene may not fully capture the nuanced composition and dynamics of Notch signaling, as well as the activation of its downstream targets. There is a need for innovative genetic tools that can more accurately represent Notch responsiveness within the hepatic biliary system. Given that *her2*, *her15.1*, and *her15.2* belong to the *hes5* gene family and exhibit similar expression profiles in the hepatic duct as revealed by single-cell profiling (Supplementary Fig. 1), we developed 3’ knock-in zebrafish models to reflect the natural *hes5* gene-mediated Notch activity (Fig. 2A). Fluorescent imaging revealed that the *her2* knock-in signals are prominently localized to the fibroblast-like IHD in 6 dpf larvae (Fig. 2B and B’), with little to no signals found in intermediate and major duct (Fig. 2C – E, Supplementary Fig. 5A - D). This expression pattern persists as the fish mature into juveniles and adults (Fig. 2C – E and Supplementary Fig. 5A - D).

**Fig. 2.**
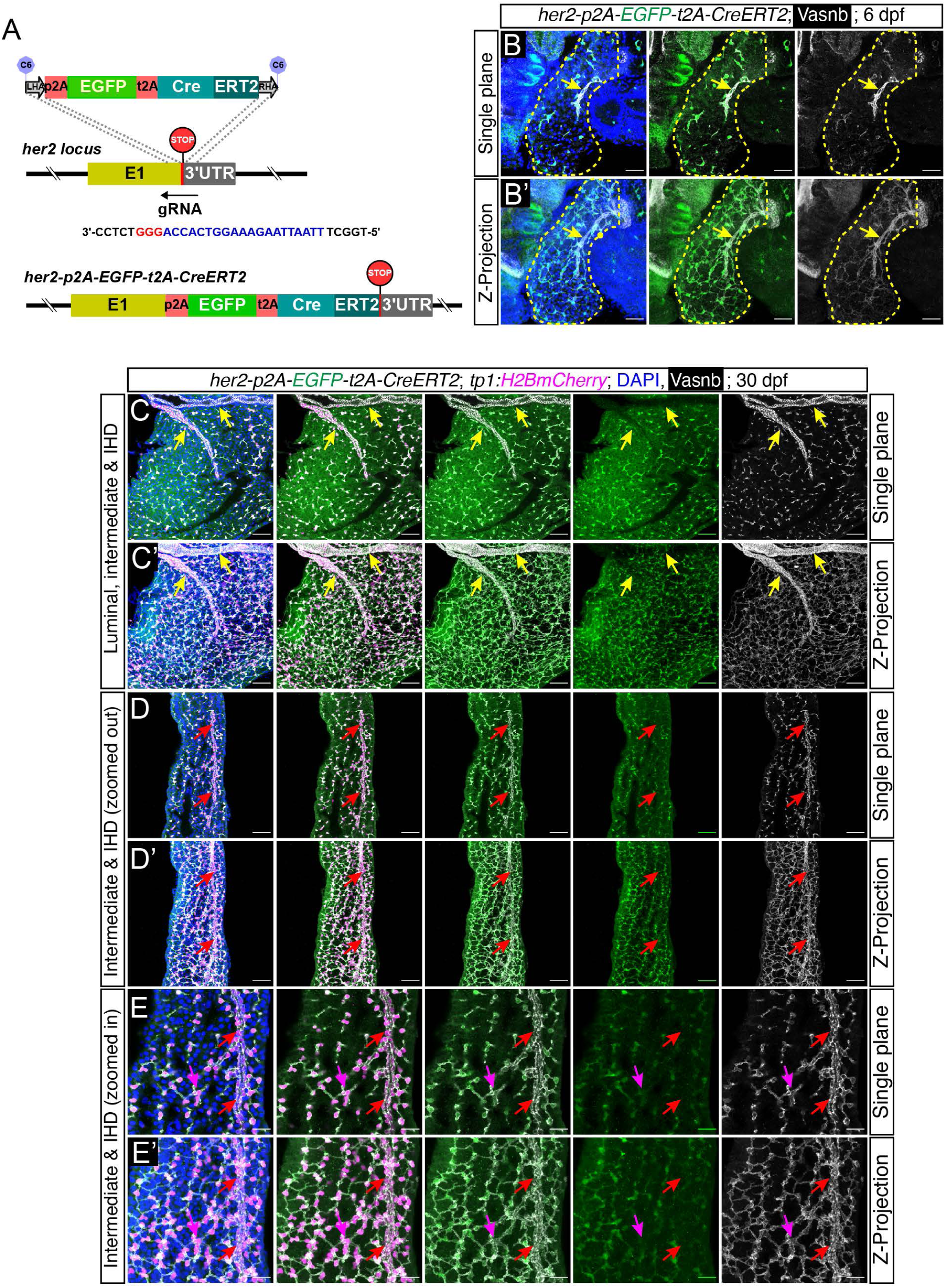
The design and characterization of a knock-in line at the *her2* locus. (A) The design of donor dsDNA templates and gRNA sequence for the construction of a knock-in line at the *her2* locus. The nucleotide sequence in blue indicates gRNA, whereas the nucleotide sequence in red indicates the PAM sequence. (B) Representative single-plane (B) and Z-projection (B’) <u>confocal images</u> showing the *her2* labeling in hepatic biliary ductal tree in the larval zebrafish liver. The yellow dashed line indicates the liver, and the yellow arrow indicates the large luminal duct that forms the stem of the ductal tree. (C-E) Representative single-plane (C,D,E) or Z-projection (C’,D’,E’) confocal images for the visualization of *her2* expression pattern in different hepatic regions using juvenile *TgKI(her2-p2A-EGFP-t2A-CreERT2)* homozygous knock-in reporter line. The arrows indicate the ducts with different morphology and spatial distribution, i.e., luminal duct (yellow), intermediate duct (red), and IHD (magenta). Scale bars, 100 μm (B-D) and 40 μm (E).

Employing a similar strategy, we successfully generated the *her6* and *her15.1*/*her15.2* 3’ knock-in zebrafish models. The *her6*-driven fluorescence was observed primarily within the IHDs, with no detectable signal in the EHDs and luminal duct inside the liver (Supplementary Fig. 3). Interestingly, in the transitional zones with intermediate ducts, adjacent to the IHDs, the green fluorescence persisted, but largely vanished in major ducts. This distinct gradient continuum of expression is maintained throughout development, from the juvenile to the adult stages (Supplementary Fig. 4E – H). The *her15.1* and *her15.2* genes exhibit identical coding and flanking sequences at their 3’ ends, and the knock-in is located within one of these nearby loci. In the 6 dpf *her15.1*/*her15.2* knock-in larvae we observed no fluorescence. However, in the juvenile and adult stages, the fluorescence was confined to a subset of IHDs (Supplementary Fig. 5E - H). To facilitate classification, we categorized the hepatic ductal trees into three main groups based on the combined expression of the three *her* genes (*her2*, *her6*, and *her9*): *her2^+^*/*her6^+^*/*her9^+^*triple positive ductal cells, which correspond to the classical *tp1*^+^Notch-responsive IHD cells; *her2*^-^/*her6*^-^/*her9*^+^ luminal ductal cells; and *her2*^-^/*her6^dim^*/*her9*^+^ intermediate ductal cells, which are residing in between the aforementioned ductal cell types.

To evaluate the responsiveness of the various *her* genes to Notch and BMP signaling pathways, we subjected homozygous knock-in zebrafish larvae to treatments with a Notch signaling inhibitor (LY411575) and a BMP inhibitor (DMH-1). Fluorescent imaging revealed a modest reduction in fluorescence intensity in *her9* knock-in larvae following exposure to a low dose of LY411575 (Supplementary Fig. 6A - E). Notably, a synergistic interaction between LY411575 and DMH-1 was observed, leading to a pronounced attenuation of *her9*-driven fluorescence. Strikingly, treatment with a low dose of LY411575 nearly eliminated the fluorescence driven by *her2* and *her6* (Supplementary Fig. 6F - M). In contrast, treatment with DMH-1 alone did not significantly affect the fluorescence intensity of any *her* gene. These findings indicate that *her2* and *her6* expression is predominantly regulated by Notch signaling, whereas *her9* expression is influenced by both Notch and BMP pathways.

### Re-analysis of public single-cell transcriptomics data with *tp1* lineage tracing reveals novel ductal markers and insights into hepatic duct remodeling

The Morrison *et al.* scRNA-seq dataset, which was aligned to an earlier version of the reference (GRCz10/danRer10), may have resulted in the omission of key genes due to inconsistencies in gene nomenclature. To identify additional ductal markers with the benefit of refined gene annotations, we re-analyzed Wolfram Goessling lab’s single-cell dataset^20^. This dataset employed the *tp1*-driven lineage tracing approach to enrich cells originating from the *tp1*^+^ lineage at various times points during liver regeneration (Fig. 3A). In addition to the cell types which the researchers demonstrated, we identified a distinct cluster representing luminal duct at the corner close to the IHD on the Uniform Manifold Approximation and Projection (UMAP) plot. This cluster exhibits a unique expression profile of *her9* along with genes encoding multiple structural proteins, such as *krt4*, *agr2*, *cdh17*, *cldnh*, and *cldn15la* (Fig. 3B), and indicates a potential conversion from the *tp1^+^*to *krt4^+^* ducts.

**Fig. 3.**
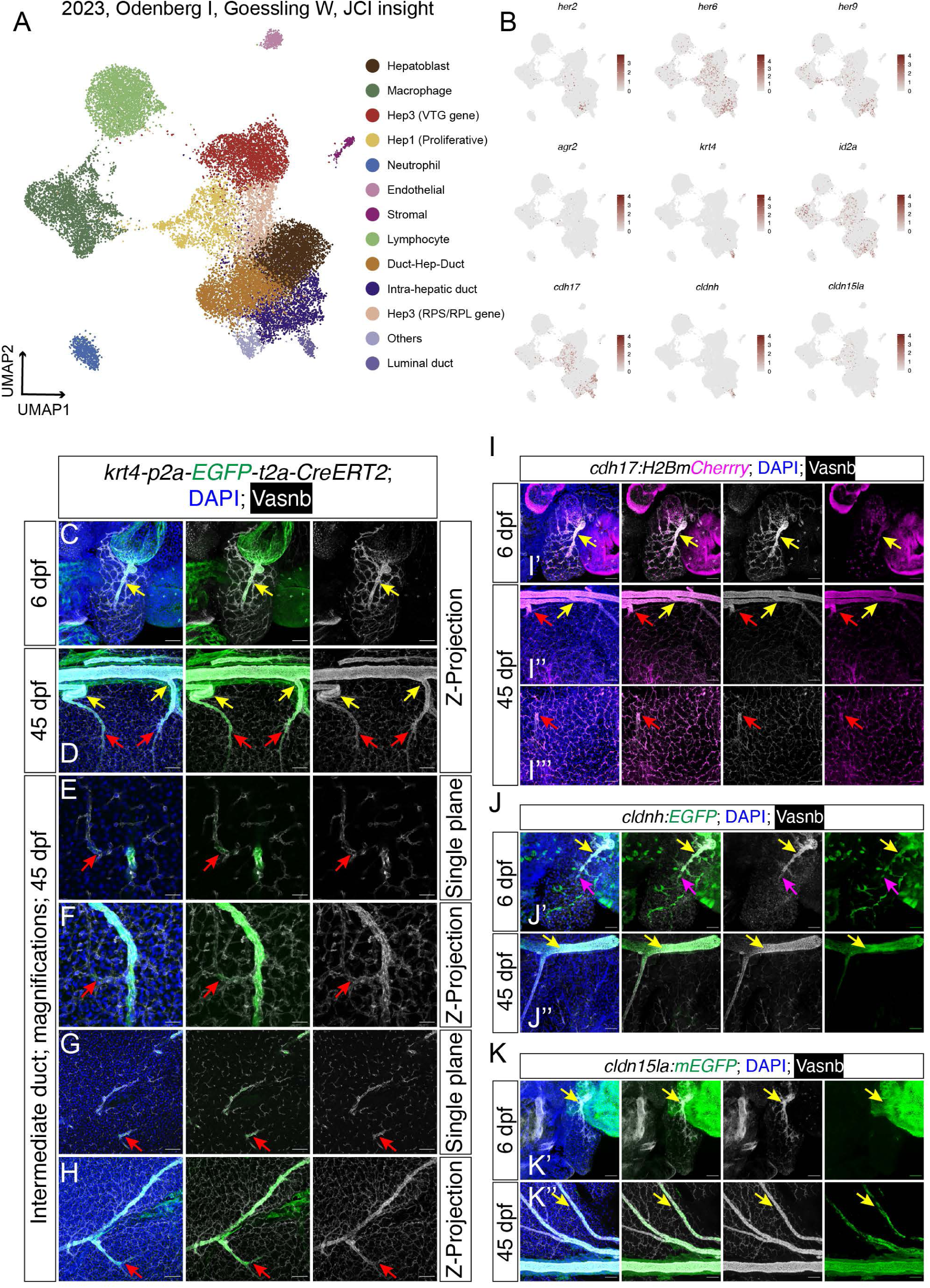
Re-analyzing scRNA-seq of *tp1* lineage tracing show ductal heterogeneity and plasticity. (A) UMAP plot showing the cell clusters captured from various timepoints in the liver regeneration experiments done by Oderberg *et al*. (B) The Feature-plot displaying the ductal marker genes highly expressed in the *krt4*^+^ ductal cluster, indicating potential *tp1*^+^-to- *krt4^+^* ductal cell conversion. (C-H) Representative single-plane or Z-projection confocal images showing the *krt4* labeling in various locations of the hepatic biliary ductal tree. (I) Representative confocal images showing the *cdh17:H2BmCherry* transgenic labeling in larval and juvenile liver ducts. (J) Representative confocal images showing the *cldnh:EGFP* transgenic labeling in larval and juvenile liver ducts. (K) Representative confocal images showing the *cldn15la:mEGFP* transgenic labeling in larval and juvenile liver ducts. The arrows point to the ducts with different morphology and spatial distribution, i.e., luminal duct (yellow) and intermediate duct (red) and IHD (magenta).

Our earlier research in zebrafish pancreas revealed that *krt4*^+^ ductal cells constituted a previously unrecognized cell type, which corresponded to the luminal ducts in both the extra-pancreatic region and within the pancreas^22^. To validate the presence of *krt4*^+^ ducts in the liver, we utilized *krt4* knock-in reporter^23^. Fluorescent imaging indicated a strikingly similar expression pattern to that observed in the pancreas, with intense green fluorescence in large luminal ducts and diminished signals in intermediate ducts, while no fluorescence was detected in the IHDs (Fig. 3 C - H).

We further confirmed the expression of *cdh17*, *cldnh*, *cldn15la*, and *agr2* using transgenic reporters. Among these, *cdh17* displayed pan-ductal expression, whereas *cldnh*, *cldn15la*, and *agr2* were enriched in luminal and intermediate hepatic ducts (Fig. 3I – K and Supplementary Fig. 7A - C). Collectively, by integrating single-cell transcriptomics with multiple knock-in and transgenic lines, we have delineated ductal heterogeneity based on their distinct molecular signatures and spatial distribution. Moreover, the unexpected finding of *krt4*^+^ luminal ducts through *tp1^+^* lineage tracing prompts us to hypothesize that the *krt4*^+^ ducts may originate from the *tp1*^+^ ducts during hepatic duct remodeling.

### Lineage tracing reveals ductal cell plasticity in static conditions and in response to IHD loss

To delve deeper into the plasticity of ductal cells, we employed a lineage tracing approach to genetically track the fate of ductal cells (Fig. 4A). Initially, we utilized the *Tg(tp1:CreERT2);Tg(ubb:CSHm)* to label the *tp1*-expressing ducts at 2-3 dpf by treating the larvae with 4-hydroxytamoxifen (4-OHT). The Cre-mediated recombination resulted in the expression of nuclear mCherry, driven by a ubiquitin promoter. Subsequently, liver samples were harvested at 35 dpf juvenile stage. Immunofluorescent staining revealed that mCherry^+^ cells were distributed in both intra-hepatic and luminal ducts (Fig. 4B and C), corroborating Goessling’s scRNA-seq findings (Fig. 3A and B). However, due to the random integration nature of the *tp1* transgenics, we cannot rule out the potential for the *tp1* transgene to be integrated into an open chromatin region with hyperactive transcriptional activity, thus leading to the erroneous labeling of *tp1^+^* cells in the *krt4*^+^ duct. To overcome this issue, we performed experiments *her2-CreERT2* knock-in and confirmed that both luminal and intermediate ducts could be genetically fate-mapped by Notch-responsive IHD labelling at an embryonic stage in static conditions (Fig. 4D and E). Collectively, these results, in conjunction with our previous pancreas research, indicate that ductal remodeling is a universal biological process that also facilitates the reorganization and maturation of fully functional ductal architecture in the hepato-pancreatico-biliary system.

**Fig. 4.**
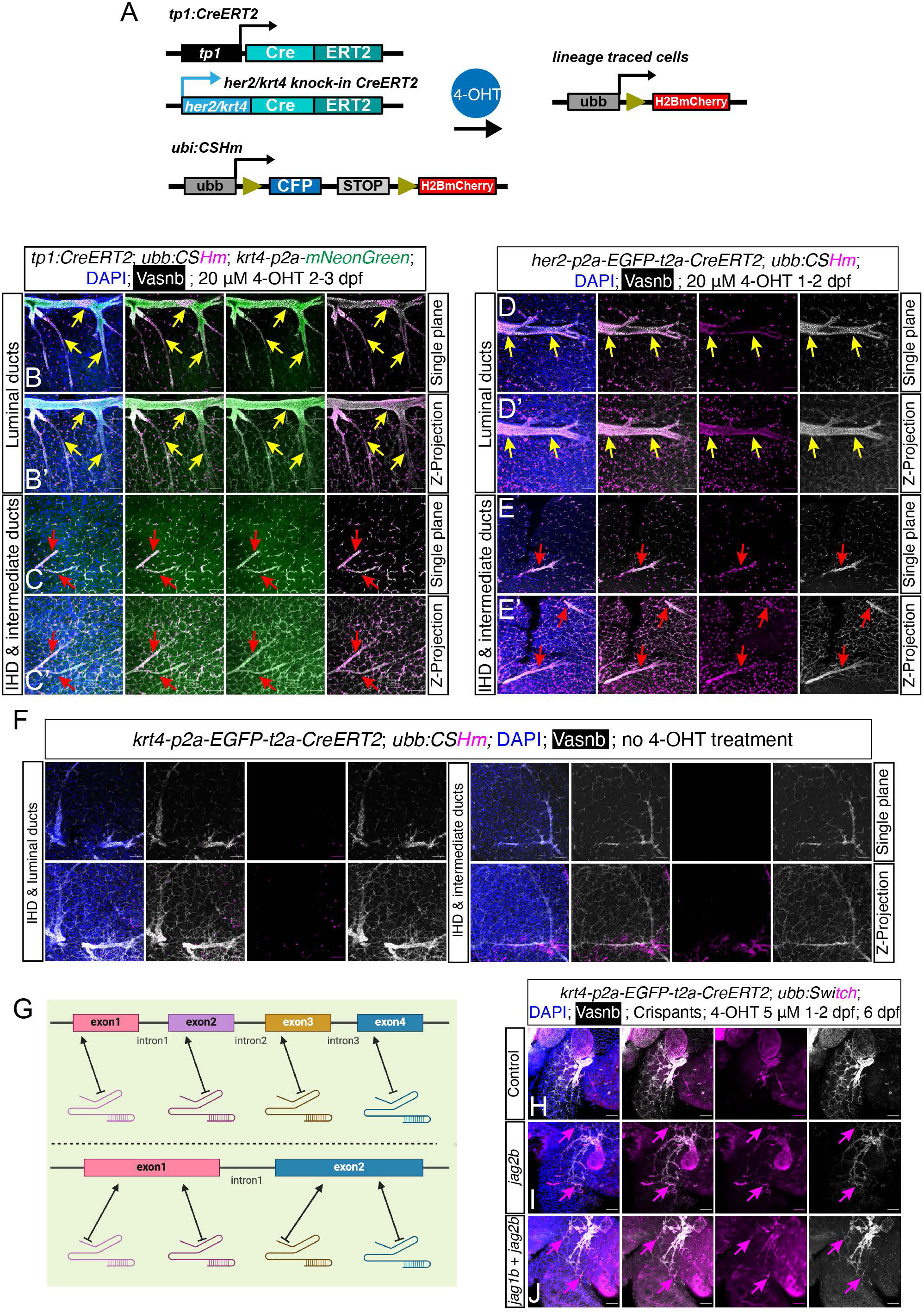
Spatiotemporal-controlled lineage tracing indicating hepatic duct cell plasticity. (A) Workflow of the Cre/loxP system used for spatiotemporal lineage tracing using different CreERT2 lines for the labeling of BECs in IHD or luminal ducts. Fish carrying CreERT2 are crossed to fish carrying the *ubb:CSHm* transgene. The 4-OHT is administered from 1 to 2 dpf or 2 to 3 dpf, and displayed as 35 dpf juvenile fish in B-F. (B-C) Single-plane (B and C) or Z-projection (B’ and C’) confocal images showing mCherry^+^ lineage traced cells in the main duct proximal to the hilum, intermediate duct and IHD in the periphery of the ductal tree using *Tg(tp1:CreERT2)* zebrafish. The ductal tree is marked by *TgKI(krt4-p2A-mNeonGreen)* and Vasnb staining. Scale bars, 100 μm. (D-E) Single-plane (D and E) or Z-projection (D’ and E’) confocal images showing mCherry^+^ lineage traced cells in the main luminal duct proximal to the hilum, intermediate duct and IHD in the periphery of the ductal tree using *TgKI(her2-p2A-EGFP-t2A-CreERT2)* zebrafish. Scale bars, 100 μm. (F) Single-plane and Z-projection confocal images showing the control experiment of lineage tracing without 4-OHT using the *krt4* knock-in CreERT2 line. Scale bars, 100 μm. (G) Schematics of CRISPR-Cas9 mediated mutagenesis using gRNAs targeting different exonic regions in the candidate genes *jag1b* (upper panel) and *jag2b* (lower panel)^60^. (H-J) Z-projection confocal images showing the *krt4* lineage-traced cells at 6 dpf fish with the scramble, *jag2b* single Crispant and *jag1b*+*jag2b* double Crispant group, respectively. Scale bar, 40 μm. The arrows point to the ducts with different morphology and spatial distribution, i.e., luminal duct (yellow) and intermediate duct (red) and IHD (magenta).

A recent study indicated that the EHD may possess the ability to undergo cell fate conversion and become IHD under extreme IHD loss conditions^26^. However, evidence from direct lineage tracing studies focusing on the EHD is currently limited. In this study, we integrated the *krt4-CreERT2* knock-in system to genetically label the luminal duct, primarily the EHD, during the embryonic stage. In the control experiments, the absence of 4-OHT treatment resulted in no mCherry fluorescent labeling of luminal ductal cells (Fig. 4F). We administered a series of Cas9-sgRNA ribonucleoprotein (RNP) complexes targeting *jag2b* alone or *jag1b* and *jag2b* in combination to induce CRISPR-Cas9-mediated mutations that simulate the loss of IHD, akin to the pathological features observed in Alagille syndrome (Fig. 4G, Supplementary Fig. 7D -E). In the 4-OHT treated control group, only the luminal ducts and cells immediately next to it in the hilum of the liver exhibited mCherry fluorescent labeling from the *ubb:Switch* responder line (Fig. 4H). In the *jag2b* Crispant group, we observed a reduction in the number of IHD cells, with a subset displaying a fibroblast-like morphology and labeled with mCherry (Fig. 4I). In the *jag1b*/*jag2b* double Crispant model, there was a significant loss of IHD cells, with only sporadic mCherry-labeled fibroblast-like ductal cells remaining in areas adjacent to the distorted luminal ductal tree (Fig. 4J). These findings underscored the plasticity of ductal cells in both homeostatic conditions and in response to a severely compromised ductal tree, where the luminal ducts can remodel to fibroblast-looking ductal cells.

### Single-cell transcriptomics unveils ductal heterogeneity in juvenile hepatic duct

The inherent limitations of ductal cell sorting methods have historically hindered the enrichment of a comprehensive array of hepatic ductal cells in the prior single-cell datasets. To address this challenge, we employed a dual transgenic system, *TgKI(her9-p2A-EGFP-t2A-CreERT2); Tg(ubb:CSHm)*, with 4-OHT treatment at 1-2 dpf to label pan-hepatic ductal lineages. At the 30 dpf juvenile stage, we isolated the liver and utilized FACS to capture the mCherry^+^ cells. After quality control (QC), a total of 4759 ductal cells were included in our analysis. The UMAP plot revealed two major clusters with distinct expression patterns of *krt4* (Fig. 5A and B). The *krt4*^+^ cluster was further resolved into two sub-clusters based on the expression levels of *cldn15la*, *cuzd1.2*, *agr2*, and *hpda* (Fig. 5B). Conversely, the *krt4*^-^ cluster exhibited specific expression of *her2*, *her15.1*, and *her15.2* (Fig. 5B), indicating IHD characteristics. Notably, this cluster also showed a marked expression of *notch3*, suggesting that its Notch responsiveness could be primarily mediated through the Notch3 receptor. This cluster can be further categorized into three major sub-clusters, with the Duct2 subtype characterized by a general downregulation of *her* genes and other ductal markers, alongside an upregulation of cell cycle-related genes (pcna and *top2a*), implying a proliferative state within the IHD. It is also noteworthy that both *her6* and *her9* were expressed across all ductal cell types, with *her6* showing a downregulation in *krt4^+^* clusters, though not to the point of absence (Fig. 5B and C). Pseudotime analysis revealed that the Duct2 subtype serves as the starting point in the trajectory, with a linear progression observed from the IHD towards the *krt4*^+^ duct under homeostatic conditions (Supplementary Fig. 8A). PROGENy pathway analysis highlighted a significant upregulation of the EGFR and MAPK pathways within the *krt4*^+^ ducts, particularly in the Duct5 subtype (Supplementary Fig. 8B). Additionally, pathways such as estrogen, PI3K, Trail, and VEGF were found to be relatively upregulated in *krt4*^+^ ducts. In comparison, the IHD sub-clusters exhibited enriched pathways related to hypoxia, JAK-STAT, WNT, TNFa, NFkB, and TGFb signaling^27^. SCENIC regulon analysis identified consistent activity and expression patterns in genes such as *atf3*, *nr4a1*, *crema*, and *tead1a* within the *krt4*^+^ ducts (Supplementary Fig. 8C). Furthermore, Gene Ontology (GO) enrichment analysis of cluster-specific marker genes indicated distinct biological functions for some of the ductal subtypes: Duct1 in stemness, Duct 3 in cell differentiation, Duct 2 in transmembrane receptor tyrosine kinase pathway, Duct3 also in epithelial morphogenesis and cell junction organization, Duct4 and 5 in ribosomal assembly and protein translation (Supplementary Fig. 8D).

**Fig. 5.**
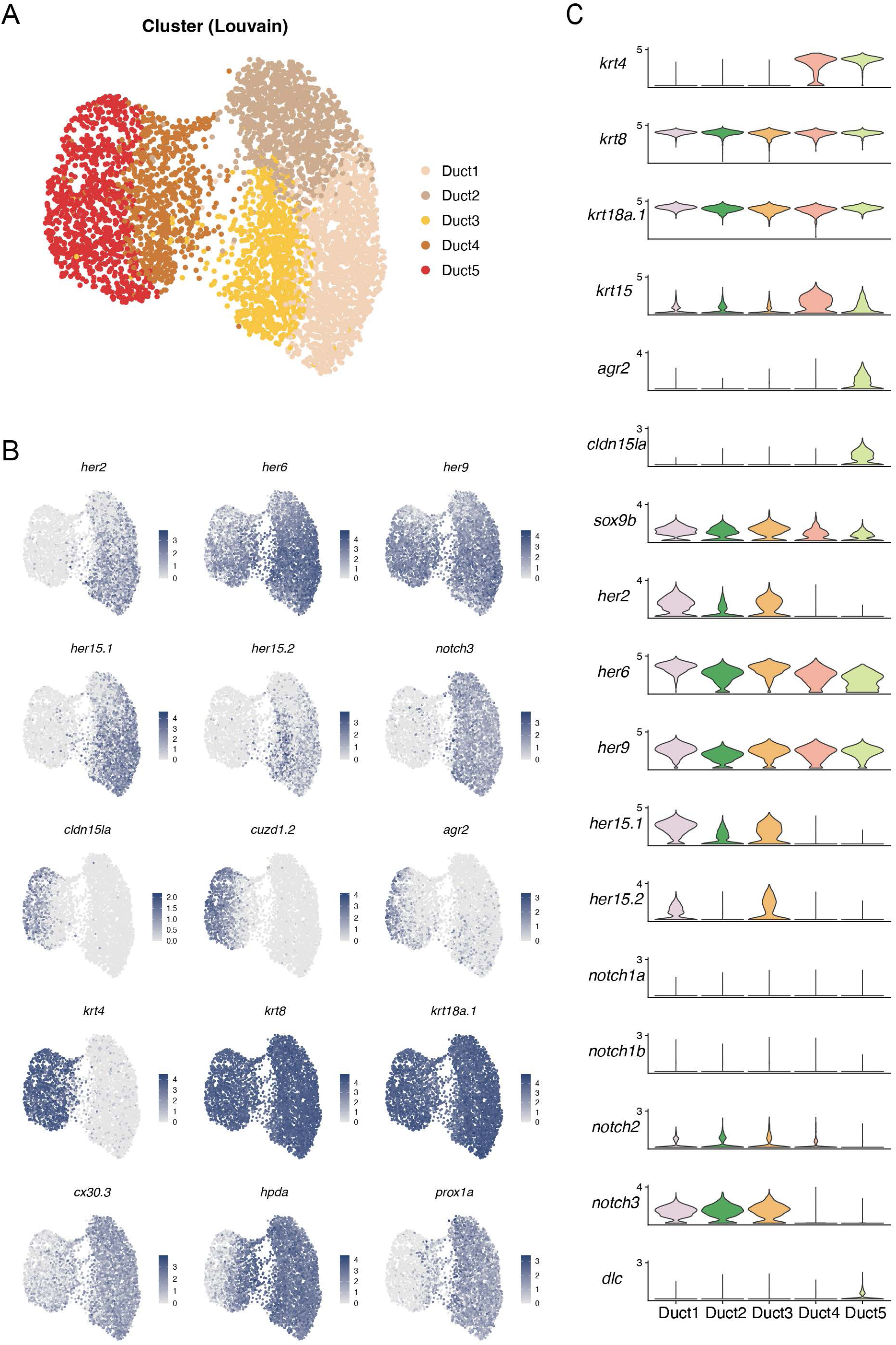
*In silico* analyses of single-cell transcriptomics showing the cell clustering in juvenile developing liver. (A) UMAP plot showing the cell clusters derived from *her9* origin in juvenile fish liver. (B) The Feature-plot showing the ductal marker genes for the *her9* lineage ductal cells. (C) The stacked violin-plots demonstrating the expression level of ductal marker genes among 5 cell clusters.

AUCell analysis, leveraging MSigDB Hallmark gene sets, revealed distinct characteristics across various ductal clusters (Supplementary Fig. 8E). The Duct1-3 exhibited heightened activity in Notch and Wnt-β-catenin signaling pathways, indicative of their developmental roles in stem or progenitor cell state maintenance^27^. In contrast, the Duct5 displayed enhanced activity in glycolysis and fatty acid metabolism, suggesting a metabolic adaptation after cell fate conversion. Both Duct1 and Duct4/5 exhibited elevated levels of p53 and mTORC1 signaling, which are pivotal to prime the cells for transdifferentiation. In our comparative assessment of nuclear protein abundance among the clusters, the pairs of Duct1/2/3 did not exhibit any single gene predominantly expressed in any cluster (Supplementary Fig. 8F). However, Duct1 was distinguished by its relatively high expression of *her2*, *her15.1*, and *her9*, while Duct3 showed a modest upregulation of *yap1* and *tead1a*. In the pairs of Duct3/4/5 (over the progression from the IHD towards the *krt4*^+^ duct), Duct3 was marked by higher expression of *her2*, *her15.1*/*her15.2*, and *prox1a* (Fig. 5B), whereas Duct5 was characterized by a significant abundance of *cdx1a* and *erg2a/b*. These transcriptional shifts in Duct3-Duct4/5 suggest profound changes occurring in response to the attenuation of Notch signaling, in concordance with previous findings^12^.

### Single-cell transcriptomics in juvenile fish at 2-day post hepatocyte ablation reveals dynamic transcriptional shifts in ductal cells

Next, we investigated the transcriptional alterations in cells of the *her9* lineage at 2 days post-hepatocyte ablation (2 dpa). We used the triple transgenic juvenile fish, *TgKI(her9-p2A-EGFP-t2A-CreERT2); Tg(ubb:CSHm); Tg(fabp10a:CFP-NTR*), and applied 4-OHT from 1-2 dpf. Hepatocyte ablation was executed at 27 dpf by treating the zebrafish with metronidazole (MTZ) for 1.5 days. Following ablation, the fish were permitted to recover for 48 hours after the removal of MTZ. We isolated mCherry^+^ cells from the liver’s single-cell suspension by FACS, yielding 3079 cells that passed QC. The UMAP plots displayed three main ductal clusters and four clusters exhibiting hepatocyte-like characteristics (Fig. 6A). Two of the ductal clusters (Duct1 and Duct2) displayed a distinct *krt4* expression (Fig. 6E) while the remaining ductal cluster (Duct3) expressed *her2*, *her15.1*, and *her15.2*, indicative of Notch-responsive IHD cells, and all three ductal clusters uniformly expressed *sox9b* (Supplementary Fig. 9), in line with our previous single-cell findings under static conditions (Fig. 5).

**Fig. 6.**
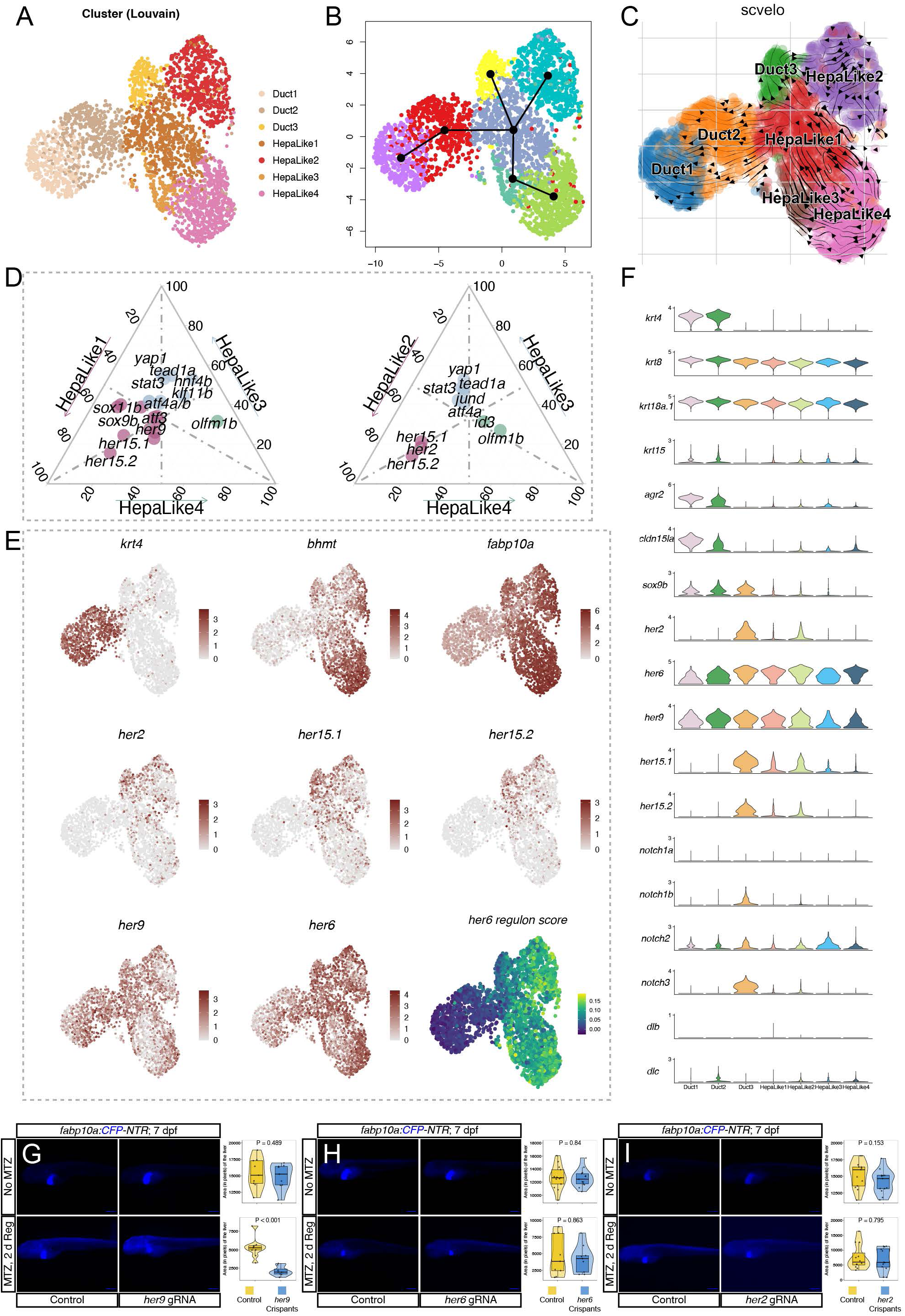
*In silico* analyses of single-cell transcriptomics showing the cell state transition during liver regeneration. (A) UMAP plot showing the cell clusters derived from *her9* origin labeled via 10 μM 4-OHT treatment 1-2 dpf, and harvested from juvenile fish liver at 2 dpa. (B) Slingshot pseudotime trajectory showing the cell transition flow with HepaLike1 as the starting point. (C) The scVelo vector showing the cell transition direction from the HepaLike1 towards other cell clusters. (D) Ternary plot shows the expression of DNA binding genes in pairs of HepaLike1/3/4 pairs of HepaLike2/3/4. (E) The Feature-plot and the UMAP of liver marker genes in the *her9* lineage, as well as the *her6* regulon activity score. (F) The stacked violin-plots demonstrating the expression level of marker genes among 7 cell clusters. (G-I) The immunofluorescent imaging and quantification results showing the regenerative liver size in *her9*, *her6* and *her2* Crispants carrying the *fabp10a:CFP-NTR* transgene.

**Fig. 7.**
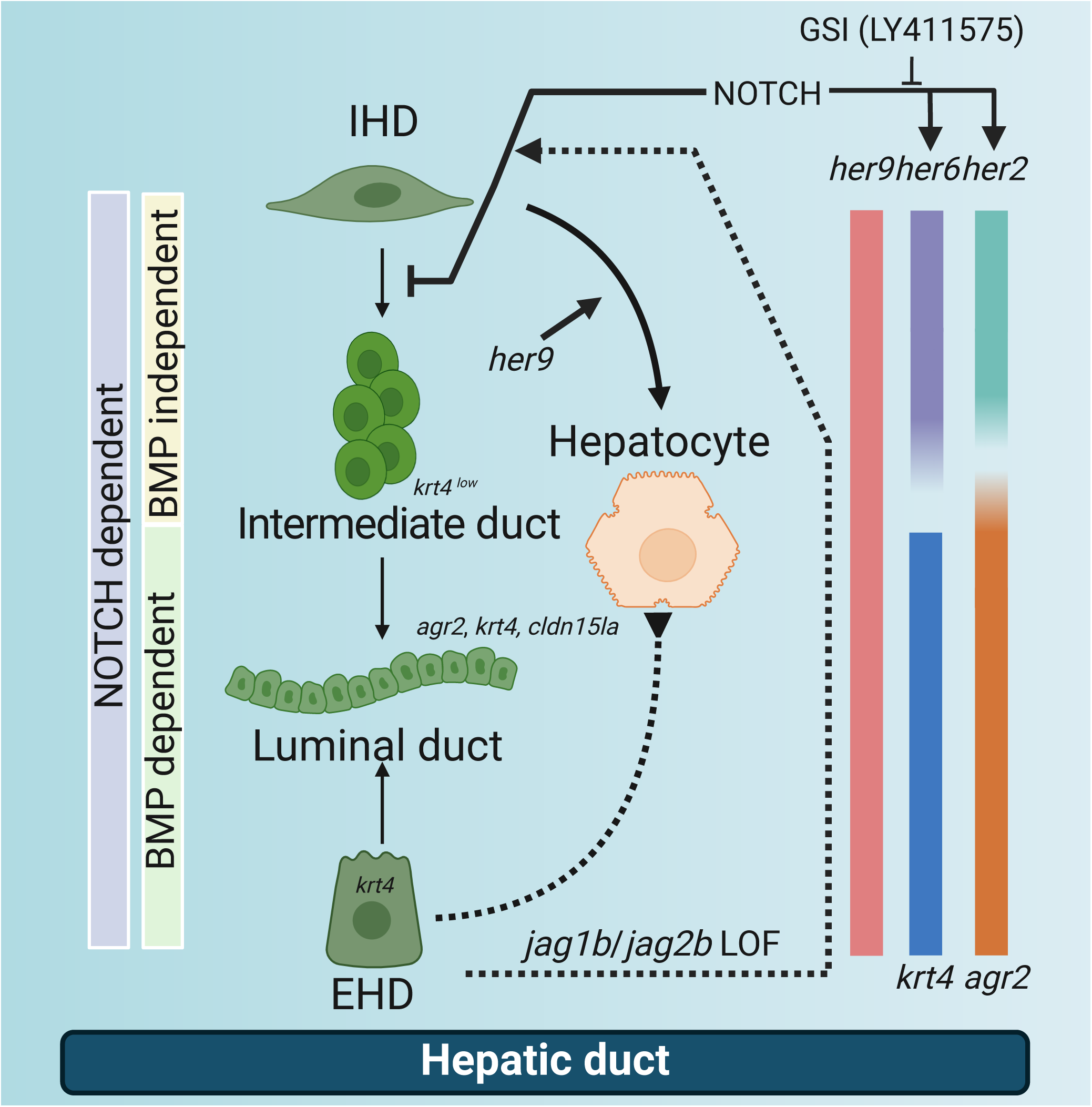
Schematic diagram. Summary of the hepatic duct heterogeneity, marker genes, cell plasticity and dynamics, and the responsiveness to signaling pathways; created by Biorender, https://www.biorender.com/.

Among the hepatocyte-like (HepaLike) clusters, HepaLike1 was centrally positioned within the embedding, whereas HepaLike2, located at the upper right corner, demonstrated a specific upregulation in cell cycle-associated genes, including *top2a*, *pcna*, *mki67*, and *ccna2* (Supplementary Fig. 9). HepaLike3 exhibited upregulation of *stat3*, *tg* (encoding thyroglobulin), and *yap1*. HepaLike4 clusters presented the most pronounced expression levels of hepatocyte markers and functional proteins, encompassing the hepatocyte maturation marker (*bhmt*), transcription factors (*hnf4a* and *hnf4b*), fatty acid binding protein (*fabp10a*), coagulation factors (*f2* and *f10*), complement protein (*c1r*), and apolipoprotein (*apoa1b*) (Fig. 6E, Supplementary Fig. 9).

Pseudotime trajectory analysis, complemented by velocyto analysis, positioned the HepaLike1 cluster as the starting point, functioning like the progenitor, which is capable of differentiating or re-differentiating into hepatocytes or various ductal cell types (Fig. 6B and C). Feature-plot and ternary plot visualizations underscored the notable expression of *her6* and *her9* within the HepaLike1 cluster (Fig. 6D - F), in the absence of Notch receptors (Supplementary Fig. 9). AUCell analysis revealed heightened activity in pathways such as mTORC1, the unfolded protein response, and the PI3K-AKT-mTOR signaling cascade in HepeLike1 cluster (Supplementary Fig. 10A). PROGENy pathway analysis further indicated the upregulation of the PI3K, WNT, and VEGF pathways in HepaLike1 (Supplementary Fig. 10B). GO enrichment analysis attributed functions associated with ribosome biogenesis, RNA splicing, mRNA processing, and protein translation to HepaLike1 (Supplementary Fig. 10C). SCENIC regulon analysis highlighted the upregulation of the *her6* regulon in the HepaLike clusters (Fig. 6E - F), importantly also in HepaLike1 that has low Notch activity (Supplementary Fig. 10). Additionally, increased regulon activity was observed in other cell types of interest, including for *atf6* and *cebpa* in HepaLike1, 2 and 4, *dnmt1* in HepaLike2 and 3, and *nr4a1* in Duct1 and 2 (Supplementary Fig. 11). Given that *her6* and *her9* are members of the *hes1* gene family, have similar DNA binding domains, and exhibit analogous expression patterns, it is plausible that the *her6* regulon is also representing activity of *her9*-expression.

To investigate the functional role of *her* genes in liver regeneration, we used CRISPR mutagenesis to generate *her2*, *her6*, and *her9* Crispants, each targeted by 4 different sgRNAs. Experimental comparisons were made between these Crispants and scramble-injected controls. Assessment of fluorescent signals from *fabp10a:CFP-NTR* after hepatocyte ablation indicated that the mutagenesis of specifically *her9* resulted in a significantly smaller liver at 2 dpa (Fig. 6G - I). This finding suggests that *her9* is crucial for the duct-derived recovery of hepatocyte function.

## Discussion

In this study, we delved into the intricacies of the hepatic ductal system, with focuses on the insights of the heterogeneity and plasticity of hepatic ductal cells during development and liver regeneration. Utilizing the state-of-the-art zebrafish models coupled with single-cell transcriptomics, we identified distinct molecular signatures, especially the differential expression pattern of the *her* family genes in distinguishing various types of hepatic ductal cells. Our findings uncover *her9* as a pan-ductal marker, responsive to liver regeneration. We also demonstrate novel ductal markers and the dynamic remodeling process during development and injury response, highlighting the remarkable adaptability of hepatic ductal cells in maintaining cellular composition and tissue homeostasis. This discovery not only enhances our understanding of liver regeneration mechanisms in the model organism but also provides new perspectives for therapeutic strategies in severe liver diseases.

The heterogeneity of liver ductal cells has been an unexplored area and previous literature were mainly focusing on the *tp1*^+^ IHD cells as these cells have been recognized as the reservoir for regenerated hepatocytes^3, 4^. However, such cell type only forms the ductal network without an organized luminal structure, indicating that they are unable to guide the bile acid to move towards the intestine and unrecognized luminal duct cell types should exist^26, 28^. One recent study by Morrison *et al.*, first introduced the existence of hepatic ductal cell types (i.e., the *agr2*^+^ duct) other than *tp1*^+^ IHD by conducting single-cell transcriptomics on adult zebrafish livers^19^. However, new efforts are still needed as their approach was unbiased sequencing without the enrichment of liver biliary cells, which could underestimate the heterogeneity of sparse ductal cell types and cell states. Also, the use of adult zebrafish may not fully represent the complexity of developing hepatic duct, especially the ductal cells undergoing cell fate conversion and rearrangement towards mature ductal structures. Furthermore, this dataset relied on the danRer10 reference genome, which could lead to the omission of key genes when combined with the new dataset leveraging danRer11 reference genome due to gene nomenclature inconsistencies. Additionally, the absence of appropriate tools for visualizing the hepatic ductal tree precluded the biological validation of the unknown hepatic ductal cell types.

Here, in our study, we first used systematic approaches by combining the former single-cell transcriptomics datasets and anti-Vasnb immunofluorescent staining showing the whole hepatic ductal structure^21^. We identified that *her9* is a proficient pan-hepatic duct marker. With the help of the *her9* knock-in zebrafish line as well as the lineage tracing strategy to capture *her9*^+^ cell lineage, we could maximize the enrichment of liver ductal cells for the following single-cell transcriptomic studies. Additionally, we constructed multiple knock-in models based on different *her* family members. Using the combination of *her* family gene expression patterns (*her2*, *her6* and *her9*), we defined liver ductal cell types at the juvenile stage, which is a critical period of time for organ reconstruction^29^. Intriguingly, we introduced a concept of the intermediate duct, which is in between the luminal duct and fibroblast-looking IHD in terms of cell morphology, spatial distribution, and *her* family gene expression profiles. Furthermore, employing various transgenic reporter lines, we successfully validated several genes encoding structural proteins that serve as markers for liver ductal cells. Collectively, our study systematically describes the composition, morphology, and spatial orientation of liver ductal cells in zebrafish, laying an essential foundation for subsequent cell dynamic analyses of the hepatic ductal cell plasticity.

Our recent study of the zebrafish pancreas was the first to identify *krt4*^+^ ductal cells. These luminal ducts arise from *tp1*^+^ intra-pancreatic ductal cells through a remodeling process taken place during the juvenile stage. Similarly, we found *krt4*^+^ ductal cells that constitute the EHD and luminal ducts within the liver^21, 26, 30, 31^. To extend the previous finding and ascertain the remodeling is an universal process in luminal duct formation, we employed *Tg(tp1:CreERT2)* and *TgKI(her2-p2A-EGFP-t2A-CreERT2)* for the specific labelling of IHD using genetic fate mapping. Notably, the *her2* knock-in circumvents potential mislabeling issues which could happen in transgenic lines due to random insertion and unpredictable transcriptional activation, thus providing a more precise tool in delineating cell lineages^23^, but may not capture all ductal cell types/states. Our long-term lineage tracing results corroborated the hypothesis that intra-hepatic luminal ducts originate from the fibroblast-looking IHD, a process accompanied by a decrease in the expression of *her2* and *her6* genes, which are also confirmed to be regulated by Notch signaling^32^. This finding is consistent with our immunofluorescent staining demonstrating the intermediate ducts are characterized by cell aggregation and loss of apical-basal polarity^32^. Therefore, the remodeling of Notch-responsive ducts is a crucial and universal step in the formation of luminal ducts in both liver and pancreas during development. Interestingly, one recent study by Zhao *et al*, showed that inhibition of the Notch signaling can lead to the aggregation of ductal cells in the liver, and such process can be reversed to some extent when Notch inhibition is released^32^. These observations further underscore the essential role of Notch signaling in triggering cell coalescence and remodeling of IHDs during development.

To elucidate the transcriptional landscape of liver duct during growth, we selected zebrafish at the juvenile stage for further investigation. We employed the *her9* lineage tracing approach to capture a broad and substantial number of liver ductal cells. We found two major subtypes and five cellular states in the hepatic duct. The expression profiles of *krt4* and the *hes5* family genes, *her2*, *her15.1*, and *her15.2*, allowed us to distinguish two main types of ductal cells. Furthermore, we corroborated the expression of additional liver duct-related marker genes. The global gene expression profiles revealed the existence of transitional states within both subtypes. Intriguingly, although *her6* was expressed across all liver ductal cell states, its expression was notably lower in *krt4*^+^ ductal cells compared to *krt4*^-^ IHDs (too low for the *her6* knock-in line to drive detectable EGFP in *krt4*^+^ ducts). Employing hallmark gene sets archived in MSigDB and the expression profiles, we inferred that *tp1*^+^ ductal cells might activate the Notch signaling mediated by the Notch3 receptor. This finding adds a layer of complexity to our understanding of Notch signaling in ductal cells and hints at the potential involvement of alternative Notch receptors in mediating cellular transitions and phenotypes within the liver duct.

To further explore the plasticity of ductal cells and their reactivity upon severe liver injury, we again employed the *her9* knock-in lineage tracing approach in the juvenile chemical-induced liver injury fish model at 30 dpf. Our analysis revealed the presence of three ductal cell subtypes and four hepatocyte-like cell (HepaLike) clusters. The *krt4^+^* ductal cells were consistent with those observed in the aforementioned basal state, while the IHD cells only formed a single cluster. Notably, our pseudotime and velocyto analyses indicated that the HepaLike1 cells serve as starting point for the multiple cellular trajectories, capable of differentiating or re-differentiating into *krt4^+^*luminal ducts, *tp1*^+^ IHDs, and various HepaLike subtypes. This strongly suggested that the HepaLike1 cells may possess a multipotent progenitor-like feature akin to hepatoblasts^13–16, 33–35^. Additionally, HepaLike1 cells highly express *her6* and *her9*, while the other three HepaLike clusters express markers indicative of hepatocyte maturation (*bhmt*) and diverse liver functions, such as coagulation, complement, and apolipoprotein^20, 36^. In particular HepaLike2 upregulated cell cycle-related genes, indicating an active proliferative state to compensate for the lost hepatocytes. HepaLike3 expresses genes associated with thyroid hormone and immune-inflammatory responses, such as *tg* and *stat3*^37^, while HepaLike4 exhibits high glycolysis and fatty acid metabolism functions, suggesting its essential role in performing normal liver functions during regeneration. It is also noteworthy that HepaLike1 cells display high activity of the PI3K-AKT-mTOR pathway, which corroborates previous findings that this pathway is indispensable for normal duct-to-hepatocyte transdifferentiation^13, 38^. Thereafter, we hypothesize that targeting the genes highly expressed in HepaLike1 could affect the liver regeneration process. Given the critical role of *her* family genes in mediating Notch signaling and cell fate determination, we utilized CRISPR-Cas9 mutagenesis methodology to mutate *her2*, *her6*, and *her9*^39^. The *her9* Crispants showed a significantly smaller liver after 2 days regeneration, indicating a marked delay in liver recovery. Thus, these results suggest that targeting the *her9* gene may be a key effector in enhancing the liver regeneration.

## Conclusion

In conclusion, our study underscores the pivotal role of *her* genes in the molecular phenotyping of hepatic ductal cells, revealing a spectrum of ductal heterogeneity that is central to our understanding of liver regeneration. The remodeling of the ductal cells illustrates a dynamic process and highlights the remarkable ductal cell plasticity.

The comprehensive single-cell profiling provides the profound insights into the characteristics of ductal cells, their dynamic transcriptional changes during the contribution to hepatocyte recovery and highlights the functional pivotal roles of *her* genes in liver regeneration. These findings collectively illuminate the complex orchestration of cellular compositions and transitions within the liver, pointing towards *her* genes as key regulators of ductal cell fates and functions.

## Materials and Methods

### Zebrafish lines and husbandry

The following published transgenic and knock-in zebrafish lines were used in this study: *Tg(Tp1bglob:H2BmCherry)^S939^* ^25^ abbreviated as *Tg(tp1:H2BmCherry)*, *Tg(Tp1bglob:eGFP)^um14Tg^*^17^ abbreviated as *Tg(tp1:EGFP)*, *Tg(EPV.Tp1-Mmu.Hbb:Cre-ERT2,cryaa:mCherry)^s959^* ^40^ abbreviated as *Tg(tp1:CreERT2)*, *TgKI(krt4-p2A-EGFP-t2A-CreERT2)^KI129^ ^23^* abbreviated as *TgKI(krt4-CreERT2)*, *TgKI(krt4-p2A-mNeonGreen)^KI127^* ^23^, *Tg(-3.5ubb:loxP-EGFP-loxP-mCherry)^cz1701^* ^41^ abbreviated as *Tg(ubb:Switch)*, *Tg(UBB:loxP-CFP-STOP-Terminator-loxP-H2B-mCherry)^jh63^*^18^ abbreviated as *Tg(ubb:CSHm)*, *Tg(fabp10a:CFP-NTR)^s891Tg^*^42^, *Tg(cldnh:GFP) ^43^*, *TgBAC(cldn15la:GFP)^pd1034Tg^*^44^.

Several lines were re-established by re-injecting the following constructs: *Tg(-2.5cadherin17:mCherry)* abbreviated as *Tg(cdh17:H2Bmcherry)* (plasmid from Dr. Bing He)^45^, and *Tg(-6.0agr2:EGFP)* abbreviated as *Tg(agr2:EGFP)* (plasmid from Prof. Sheng-Ping L. Hwang)^46^.

The *TgKI(her9-p2A-EGFP-t2A-CreERT2)^KI151^*, *TgKI(her6-p2A-EGFP-t2A-CreERT2)^KI144^*, *TgKI(her2-p2A-EGFP-t2A-CreERT2)^KI150^* and *TgKI(her15.1/her15.2-p2A-EGFP-t2A-CreERT2)^KI152^*lines were generate according to our knock-in strategy described previously^23^, using PCR amplified 5’-AmC6 modified double-stranded DNA as the donor) and the following primers and sgRNAs:

sgRNA (*her9*): TGAGATGCGCAAGTCTACCA(GGG)

sgRNA (*her6*): GGCGGCCTTGGTAGACTCCG(AGG)

sgRNA (*her2*): TTAATTAAGAAAGGTCACCA(GGG)

sgRNA (*her15.1/her15.2*): GTGTTCGAGCGTCTCTACCA(GGG)

forward primer (*her9*): GGAGCAGAAAGCAATGAGCCGGTGTGGAGACCCTGGGGAAGCGGAGCTACTAACTTCA GC

reverse primer (*her9*): ATGAAAACTTTATAAGTTCATATGAGATGCGCAAGTCTAAGCTGTGGCAGGGAAACCCT C

forward primer (*her6*): CCGTCCGTCACGTCAGACTCCGTTTGGCGGCCTTGGGGAAGCGGAGCTACTAACTTCA GC

reverse primer (*her6*): AAGAGTCTTTTAAGAGTTTTTTTTTCTCCTCGGAGTCTAAGCTGTGGCAGGGAAACCCTC

forward primer (*her2*): TCAGAAATCGCCAAGCATGGATTGTGGAGACCCTGGGGAAGCGGAGCTACTAACTTCA GC

reverse primer (*her2*): TTTATAAAAACAACACGTTTGGCTTTAATTAAGAAAGGTCAAGCTGTGGCAGGGAAACCC

forward primer (*her15.1/her15.2*): GAGCCGCGGGCGCACGCTCCGCTCTGGAGACCCTGGGGAAGCGGAGCTACTAACTTC AGC

reverse primer (*her15.1/her15.2*): TAAGAGACGGTAAACCACCATCTAGTGTTCGAGCGTCTCTAAGCTGTGGCAGGGAAACC C

All knock-in lines were confirmed to have a correct seamless integration by Sanger sequencing of both the 5’ and 3’ integration site. Zebrafish of both sexes, ranging from 3 months to 2 years of age, were utilized for breeding purposes. The larvae were reared in E3 medium, a solution composed of 5 mM NaCl, 0.17 mM KCl, 0.33 mM CaCl_2_, and 0.33 mM MgSO_4_, adjusted to pH 7.4 with sodium bicarbonate, at a constant temperature of 28.5 °C until they reached 6 days post-fertilization (dpf). Following this stage, juvenile and adult fish were kept under a 14-hour light to 10-hour dark cycle, with the temperature maintained at 28°C. The study, involving the use of zebrafish, was granted approval by the Karolinska Institutet Animal Care and Use Committee, and all procedures were carried out in compliance with local and regional ethical guidelines and regulations.

### Lineage tracing with tamoxifen-inducible Cre recombinase

We employed CreERT2 lines to conduct strict genetic fate mapping. These CreERT2 lines were bred with the *Tg(ubb:CSHm)* or *Tg(ubb:Switch)* line for lineage tracing assessments. For temporal labeling, zebrafish embryos harboring the CreERT2 construct and the *ubb:CSHm* or *ubb:Switch* transgene were exposed to a 20 μM concentration of 4-OHT (Sigma-Aldrich) from 1 to 2 dpf, unless specified otherwise. The treatment was administered in E3 medium within 24-well plates, housing 4 to 8 embryos per well, over a 24-hour period without medium replacement. This protocol aimed to achieve optimal labeling efficiency while ensuring the survival rate of the embryos was not adversely affected.

### Sample fixation for immunostaining

Prior to fixation, the zebrafish larvae were humanely euthanized using tricaine (Sigma-Aldrich) dissolved in E3 medium, followed by thorough rinsing with distilled water to remove any residual medium. Subsequently, the specimens were immersed in a fixing solution consisting of 4% formaldehyde (Sigma-Aldrich) in phosphate-buffered saline (PBS) (ThermoFisher Scientific) at 4 °C for a minimum of 24 hours to ensure adequate fixation. Afterward, the fixation process was halted by rinsing the samples with PBS three times to eliminate the formaldehyde. The skin and the crystallized yolk present in the larvae were carefully dissected away to unmask the liver for subsequent immunostaining procedures. For juvenile or adult zebrafish, we isolated the liver and proceeded with the aforementioned fixation method.

### Immunostaining

We adhered to the immunostaining procedure as detailed in the work of Liu *et al.*^47^. In summary, the samples were first soaked in a blocking solution consisting of 0.3% Triton X-100 and 4% BSA in PBS at room temperature for a minimum of one hour. This was followed by an overnight incubation with primary antibodies at 4 °C. Post incubation, the samples were subjected to a series of washes using a buffer containing 0.3% Triton X-100 in PBS, repeated at least ten times to ensure thorough cleansing. Subsequently, the samples were treated with a blocking solution containing fluorescent dye-linked secondary antibodies and the nuclear stain DAPI (ThermoFisher Scientific), and this incubation was carried out at 4 °C overnight. The next day, the samples were again washed with the washing buffer ten times to remove any unbound antibodies.

The primary antibodies utilized in this process included chicken anti-GFP (diluted 1:500, Aves Labs, catalog number GFP-1020), goat anti-tdTomato (diluted 1:500, MyBioSource, catalog number MBS448092), rabbit anti-insulin (diluted 1:100, Cambridge Research Biochemicals, a custom preparation), mouse anti-glucagon (diluted 1:50, Sigma, catalog number G2654), rat anti-somatostatin 1.1 (diluted 1:50, GeneTex, catalog number GTX39061), and rabbit anti-Vasnb (diluted 1:1000, a custom serum provided as a gift from Dr. Paolo Panza).

### Confocal imaging

Prior to confocal microscopy, the stained samples were mounted onto microscope slides using the VECTASHIELD Antifade Mounting Medium from Vector Laboratories. For visualization, we utilized the Leica TCS SP8 platform, opting for a 40× objective lens to capture clear images of the liver in larvae up to 7 days post-fertilization (dpf). For older juveniles and adults beyond 30 dpf, we switched to a 25× objective lens to accommodate their larger size. The resulting images were then processed and analyzed using the Fiji software^48^.

### Chemical ablation of hepatocytes and drug treatment

The hepatocyte ablation in zebrafish larvae was executed by immersing the larvae, which harbored the *fabp10a:CFP-NTR* transgene, in a solution of 10 mM metronidazole (MTZ, Sigma-Aldrich) that was first dissolved in DMSO (VWR) and was further diluted to the final concentration with the E3 medium enriched with 0.2 mM 1-phenyl-2-thiourea (PTU, Acros Organics), and this treatment was applied from 3 to 4.5 dpf for a duration of 36 hours. For the hepatocyte ablation in the juvenile, a similar approach was taken, but the concentration of MTZ was adjusted to 1 mM.

For the chemical treatment of the zebrafish larvae, the chemicals were dissolved in DMSO and added directly into the E3 medium, and this mixture was applied for a period of 48 hours, unless specifically indicated otherwise. The specific chemicals utilized in these treatments were 10 μM DMH-1 (Tocris, catalog number 4126) and 1 μM LY-411575 (Selleckchem, catalog number S2714).

### Juvenile zebrafish liver dissection and FACS

We commenced the liver isolation by euthanizing the juvenile fish (male or female) through a 5-minute immersion in an ice-cold bath of Hanks’ Balanced Salt Solution (HBSS), devoid of calcium and magnesium. Utilizing blunt forceps, we meticulously dissected the fish to excise the skin, kidneys, reproductive organs in females (eggs), and the gallbladder, thereby unmasked the liver. This intricate dissection was essential to preclude any contamination from *her9*^+^ cells originating from extraneous organs, and we endeavored to eliminate as much adipose tissue as feasible. Subsequently, we made incisions at the intestinal bulb and the hindgut’s anterior region and meticulously relocated the intestine, liver, and pancreas to a fresh dish.

Employing blunt dissection instruments, we separated the liver from the intestine and concurrently detached any attached spleen, known to be in proximity to the intestine. The entire liver was then transferred and submerged in 5 mL of ice-chilled HBSS. Pooling 4 to 8 samples per condition, we initiated enzymatic digestion with 600 μL of 10× TrypLE™ (ThermoFisher), enriched with 60 μL of 100× Pluronic F-68 (ThermoFisher), and agitated the mixture at 37 °C on a shaker set at 125 rpm for 60 minutes to avert tissue clumping. To enhance the digestion process, the tissue was intermittently aspirated every 5 minutes. The enzymatic reaction was quenched by the addition of 6 mL of pre-cooled 2% BSA, followed by centrifugation at 500 *g* for 5 minutes. The supernatant was aspirated, and the pellet was resuspended and washed with 300 μL of a pre-cooled solution comprising 1% BSA, 0.1% Pluronic F-68, and 0.1% DAPI. A Corning Falcon cell strainer (Corning 352235) was employed to sift out incompletely digested tissue fragments.

During the FACS process, we identified single-cell populations by their forward and side scatter characteristics. Subsequently, we executed a negative selection employing the DAPI channel to eliminate non-viable cells and cellular fragments. Ultimately, we harnessed the mCherry channel for further gating, thereby amassing all *her9-*lineage cells into a new tube, resulting in a single-cell suspension.

### CRISPR-Cas9 mediated mutagenesis

CRISPR/Cas9 mutagenesis was conducted following the protocol established by Wu *et al*.^39^, with minor modifications. The single guide RNAs (sgRNAs) were tested for knock-out efficacies using the *her9*, *her6*, *and her2* knock-in zebrafish lines. The sgRNA generated by *de novo* synthesis from Integrated DNA Technologies. The Cas9 protein was complexed with sgRNAs to form RNP complexes and microinjected into *Tg(fabp10a:CFP-NTR)* zebrafish embryos at the one-cell stage. The zebrafish larvae were analyzed for phenotypic outcomes at various developmental stages.

We employed the CHOPCHOP website (http://chopchop.cbu.uib.no/) and designated "danRer11/GRCz11" as the reference genome together with "CRISPR/Cas9" and "knock-in" modules. The tool systematically analyzed the exon regions of the gene and prioritized the sgRNAs, taking into account their efficiency scores, self-complementarity, and mismatch count. Subsequently, those sgRNAs with high predicted targeting efficacy were subjected to manual check within the ENSEMBL genome browser, found at https://www.ensembl.org/Danio_rerio/Info/Index.

The sgRNA used for the mutagenesis were:

sgRNA1 (*her9*): GGAGTATGAGATCCACTGGC(AGG)

sgRNA2 (*her9*): CGAGAATCAACGAGAGCCTT(GGG)

sgRNA3 (*her9*): TGCACTCGTTGAATCCTGCG(CGG)

sgRNA4 (*her9*): CGATTTCTCTCTACCTGCGA(GGG)

sgRNA1 (*her6*): GTTCATGCTCGCCGGAGTCG(CGG)

sgRNA2 (*her6*): AGCGAGAATCAACGAAAGCT(TGG)

sgRNA3 (*her6*): TGATAAACCCAAAACGGCTT(CGG)

sgRNA4 (*her6*): TCGGTACTTCCCAAGAACGG(TGG)

sgRNA1 (*her2*): TGCGCATGCGCGGAGCTACT(CGG)

sgRNA2 (*her2*): GCGCATCAGCGGTGTTCTTC(AGG)

sgRNA3 (*her2*): TAGTTTAGAACAGGGCTGAC(TGG)

sgRNA4 (*her2*): GGGCTGACTGGCCTTTATTT(CGG)

### Cell encapsulation, library preparation and sequencing

Droplet-based scRNA-seq was executed utilizing the Chromium Single Cell 3′ Library & Gel Bead Kit v3 and the Chromium Single Cell 3′ Chip G, both from 10× Genomics. We loaded and encapsulated approximately 6,000 to 10,000 cells, derived from each experimental condition, into an individual v3 reaction chamber. The generation of Gel Bead Emulsions (GEMs) and the preparation of the sequencing libraries were carried out as per the manufacturer’s guidelines. Within the controller, cells were dispersed into gel beads within an emulsion, facilitating cell lysis and subsequent reverse transcription. The resulting scRNA-Seq libraries were subjected to PCR amplification across 13 cycles, after which they were pooled and denatured. The libraries were then diluted in preparation for paired-end sequencing on a NextSeq 500 platform, following the manufacturer’s recommended protocols. The acquired sequencing data were aligned to the zebrafish reference genome, GRCz11, with the incorporation of the mCherry sequence to account for the fluorescent protein tag. This alignment was performed using Cell Ranger software v5.0.1, provided by 10× Genomics, which generated a gene-by-cell count matrix employing the default settings. This matrix served as the foundational dataset for downstream analysis of gene expression patterns across the various cell populations.

### Data preprocessing

The adult zebrafish liver scRNA-Seq datasets were obtained from the Gene Expression Omnibus (GEO) database, specifically under the accession numbers GSE193043 and GSE193844 (Morrison *et al.*), and GSE217839 (Oderberg *et al.*). The Unique Molecular Identifier (UMI) counts matrix was imported into the R environment and processed utilizing the Seurat R package, version 4.0.4^49^. Cells that exhibited a gene detection count below 400 or exceeding 7000 genes were flagged as low-quality or potential doublets, and subsequently excluded from further analysis. The dataset was then normalized and scaled using default parameters, which was followed by the selection of highly variable genes (HVGs). Additional filtering to eliminate doublets was conducted employing the Doubletfinder R package^50^. Subsequent to the standard principal component analysis (PCA) for dimension reduction, we selected a range of 8 to 30 principal components (PCs) based on the expected diversity of cell types, which were then applied for graph-based clustering using the Louvain method. For cell samples collected from fish without hepatocyte ablation, among a total of 14,776 cells, we used the following cell markers to exclude non-BECs and cell contaminations: *mpeg1*, *mfap4*, and *marco* (macrophage); *ela2l*, *ela3l*, and *amy2a* (pancreatic acinar cells); *col1a1b*, *pdgfra*, and *hand2* (fibroblasts and mesothelium); *apoa1b* and *apoba* (hepatocytes); *kdrl* and *cdh5* (endothelial cells); *fcer1g*, *sla2*, *gata3*, *ccl38.6*, and *mpx* (NK cells and myeloid cells); *hmx3a*, *cdx1a*, *krt5*, *pou2f3*, and *vil1* (intestinal epithelial cells). For cell samples collected from fish after hepatocyte ablation, among a total of 12,803 cells, we used similar markers but kept cells with both liver and BEC markers for the subsequent analyses. The relationships and distinctions among various cell clusters were visualized through the Uniform Manifold Approximation and Projection (UMAP) algorithm, while batch effects were corrected using the Harmony algorithm^51^. To dissect the molecular intricacies and trace the cellular trajectory from *her9*-derived duct-to-hepatocyte lineages, each cell cluster was meticulously annotated based on established and authenticated marker genes. This was followed by the creation of subsets of specific interest, which underwent a series of processing steps including normalization, scaling, dimensional reduction, batch correction, and graph-based clustering to reveal the underlying biology of the hepatic ductal cells. The lists of genes encoding DNA binding proteins were retrieved from the human protein atlas (https://www.proteinatlas.org/).

### SCENIC analysis

The SCENIC analysis was conducted utilizing the Python-based package pySCENIC, which facilitated the generation of a gene regulatory network^52, 53^. Initially, we identified and selected a subset of 3,000 genes with the highest variability and employed their raw count data as the foundational input for the analysis. Capitalizing on recent advancements in zebrafish research, we procured a comprehensive list of known transcription factors, complete with motif annotations. These resources were instrumental in crafting an adjacency matrix that depicted the interplay between transcription factors and their presumed target genes, thus refining the gene regulatory network after the pruning of targets. Subsequently, each cell within the dataset was evaluated and assigned a Regulon Activity Score (RAS) through the application of the AUCell tool. This scoring system allowed for the quantification of the activity of each regulon, which was then utilized to generate visualizations via t-distributed Stochastic Neighbor Embedding (t-SNE) and Uniform Manifold Approximation and Projection (UMAP) techniques. For subsequent analyses, we adhered to the analytical framework previously outlined by Suo and colleagues^54^. This involved the computation of a Regulon Specific Score (RSS) matrix, designed to pinpoint regulons that were specific to particular cell types or states. We further applied hierarchical clustering techniques to the regulons, culminating in the creation of a Connection Specificity Index (CSI) matrix. This matrix delineated individual regulon modules and illuminated the interactions between regulons and their respective modules. In the last step, we calculated the average RAS across all regulons within each module, generating a composite score that reflected the activity level of each regulon module. These scores were then averaged according to cell types, enabling the identification of potential linkages between the activity of regulon modules and specific cell types. The complete dataset of transcription factors and motif annotations utilized in this analysis is accessible through the Zebrafish Embryogenesis Spatiotemporal Transcriptomic Atlas (ZESTA) at https://db.cngb.org/stomics/zesta/.

### Pseudotime trajectory and in silico fate mapping using scVelo

Pseudotime analysis was executed utilizing Slingshot (version 1.8.0) packages, post the correction of batch effects^55^. Employing the default parameters, we established the Duct2 and HepaLike1 as the foundational nodes for our pseudotime computational framework. For the assessment of RNA velocity during liver regeneration, we deployed the scVelo tool^56^. The selection of cells for this analysis was predicated on their unique barcodes, and the gene expression matrix was refined to encompass solely the 3,000 genes exhibiting the highest variability. The estimation of velocity was underpinned by the computation of moments, leveraging a subset of 15 proximate neighbors and the foremost 8 principal components. The latent time, an essential parameter for our velocity calculations, was derived from cells extracted from non-injured samples, which served as the reference point. RNA velocity was subsequently inferred from these latent time scores, employing a dynamical modeling approach. The resultant velocity vectors were then superimposed onto a pre-existing UMAP representation for a visual and interpretative synthesis of cellular dynamics.

### Enrichment analysis

We identified differentially expressed genes across various clusters by employing the ’FindMarkers’ function with specific parameters set as min.pct = 0.25 and logfc.threshold = 0.25, subsequently filtered to include only those with pvalue.adj below 0.05. For gene enrichment analysis, we applied the ’enrichGO’ function from the R package clusterProfiler (version 3.18.1), using a pvalueCutoff of 0.01 and qvalueCutoff of 0.05, with OrgDb set to org.Dr.eg.db and the pAdjustMethod set to "BH"^57^. To evaluate and contrast the activity of biological pathways between different cellular states during liver regeneration, we retrieved normalized group data and computed the pathway activity scores. This was achieved using the ’gsva’ function from the GSVA R package^58^, based on the 50 hallmark gene sets from the Molecular Signatures Database (MSigDB) version 7.5.1.

To elucidate and graphically represent the activity of these pathways at the single-cell level along the transition from HepaLike1 to other cell clusters, we harnessed the hallmark gene sets from MSigDB and employed the ’AddModuleScore’ function from Seurat to calculate the pathway activity score for each respective module. Pathway activities of each cell were estimated with PROGENy using the top 2,500 genes of each transcriptional footprint^59^.

### Statistical analysis

Replicates of the experiments were conducted independently at least twice to ensure reliability. In the case of immunostaining, the procedure was repeated to validate the expression patterns across a minimum of five individual samples, with only the most representative images featured in the illustrations. For lineage tracing studies involving juvenile and adult zebrafish, the outcomes were substantiated by a minimum of two separate experiments conducted on distinct days. Quantification of cell counts in confocal microscopy images was performed manually with the assistance of Fiji software. The schematic representation of the knock-in strategy was achieved using the "IBS" software. For statistical analysis, we employed two-tailed Mann Whitney U tests for comparing the means of two groups or Kruskal-Wallis tests for the comparison of three or more groups, unless specifically indicated otherwise. Data are presented as mean values ± standard error of the mean (SEM), with P values of ≤ 0.05 deemed to indicate statistical significance. The term ’n’ denotes the count of zebrafish in each experimental group. All statistical computations and graphical representations of data were executed on the R platform, version 4.0.2, leveraging the "ggplot2" and "smplot2" packages for visualization.

## Acknowledgments

We thank Dr. Paolo Panza (Max Planck Institute for Heart and Lung Research) for antibodies; Didier Stainier (MPI-HLR), Bing He (Karolinska Institutet), Michael Parsons (UC Irvine), Nikolay Ninov (CRTD Dresden), Michel Bagnat (DUKE) and Prof. Prof. Sheng-Ping L. Hwang for sharing the fish lines and constructs; Marina Polo Gozalo from the Biomedicum flow cytometry core facility for the assistance in FACS; Sumeet Pal Singh for comments on the manuscript; the Eukaryotic Single Cell Genomics (ESCG) facility in Stockholm funded by Science for Life Laboratory, and Anja Mezger, Samaneh Masoumi and Anastasios Glaros for technical support in scRNA-seq; the Uppsala Multidisciplinary Center for Advanced Computational Science (UPPMAX) for providing data storage resources. The schematic was created by Biorender.com.

## Data availability

Raw and processed single-cell RNA-seq data (raw data and h5) will be accessible at the Gene Expression Omnibus with the accession number when the manuscript gets accepted. All data needed to evaluate the conclusions in the paper are present in the paper and/or the Supplementary Materials.

## Funding

Research in the lab of OA was supported by funding from:

The European Research Council under the Horizon 2020 research and innovation programme (grant n° 772365)

The Swedish Research Council

The Novo Nordisk Foundation

The Swedish Cancer Society

The Swedish Diabetes Foundation

The Swedish Child Diabetes Foundation Diabetes Wellness Sweden

Strategic Research Programmes at the Karolinska Institutet (SRP Diabetes and StratRegen) JM was supported by the National Natural Science Foundation of China (Grant Nos. 82400590)

## Author contributions

J Mi: conceptualization, data curation, formal analysis, investigation, visualization, methodology, and writing—original draft, review, and editing.

L Ren: investigation, visualization, methodology, and writing—review, and editing.

KC Liu: investigation, visualization, methodology and writing—review and editing.

L Buttò: investigation and writing—review and editing.

D Colquhoun: investigation.

O Andersson: conceptualization, formal analysis, supervision, funding acquisition, validation, investigation, visualization, project administration, and writing—review and editing.

