## Supplementary Figure 1-11 for "Deciphering the zebrafish hepatic duct heterogeneity and cell plasticity using lineage tracing and single-cell transcriptomics"

### Deciphering the zebrafish hepatic duct heterogeneity and cell plasticity using lineage tracing and single-cell transcriptomics

Jiarui Mi[1,2], Lipeng Ren[1,3], Ka-Cheuk Liu[1,3], Lorenzo Buttò[1,3], Daniel Colquhoun[1,3], Olov Andersson[1,3]*

1. Department of Cell and Molecular Biology, Karolinska Institutet, Sweden
2. Department of Gastroenterology, Sir Run Run Shaw Hospital, Zhejiang University School of Medicine, China
3. Department of Medical Cell Biology, Uppsala University, Sweden

*Corresponding author

Olov Andersson

Morisson *et al.* dataset

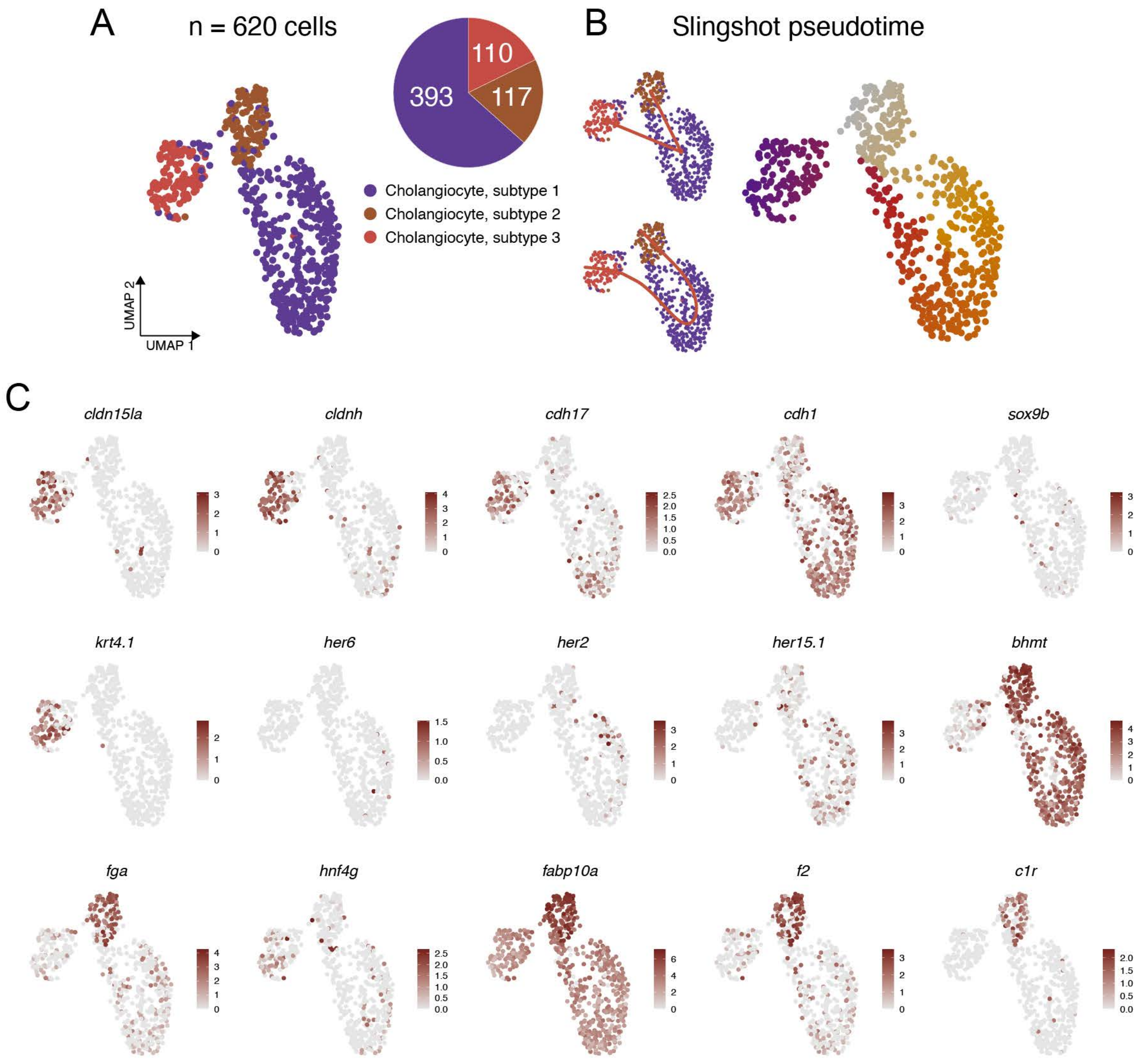

**Supplementary Fig. 1.** (A) UMAP plot showing the ductal cell clusters from the scRNA-seq done by Morrison et al. (B) The Slingshot pseudotime results of different types of ducts. (C) The feature plots showing the representative marker genes for each cluster.

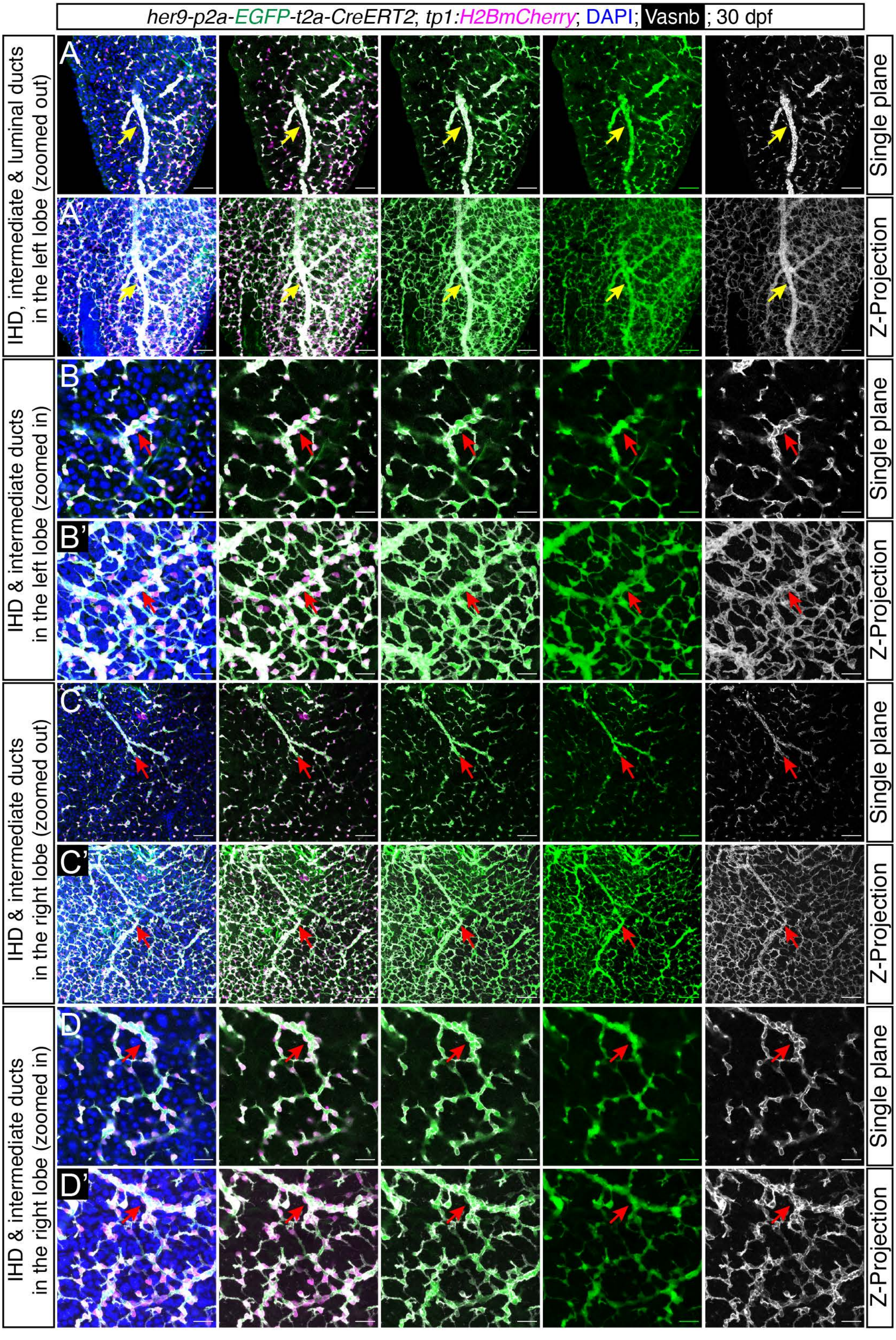

her9-p2a-EGFP-t2a-CreERT2; tp1:H2BmCherry; DAPI; Vasnb; 30 dpf
IHD, intermediate & luminal ducts in the left lobe (zoomed out)
IHD & intermediate ducts in the left lobe (zoomed in)
IHD & intermediate ducts in the right lobe (zoomed out)
IHD & intermediate ducts in the right lobe (zoomed in)
A
A'
B
B'
C
C'
D
D'
Single plane
Z-Projection

**Supplementary Fig. 2. Immunostaining showing the *her9* expression pattern in hepatic biliary duct in juvenile fish.** (A-D) Representative single-plane (A,B,C,D) or Z-projection (A’,B’,C’,D’) confocal images for the visualization of *her9* expression in juvenile zebrafish using *TgKI(her9-p2A-EGFP-t2A-CreERT2)* homozygous knock-in reporter line with magnifications shown in B,B’,D,D’. Scale bars, 200 μm (A,A’,C,C’), 40 μm (B,B’,D,D’). The arrows indicate the ducts with different morphology and spatial distribution, i.e., luminal duct (yellow) and intermediate duct (red).

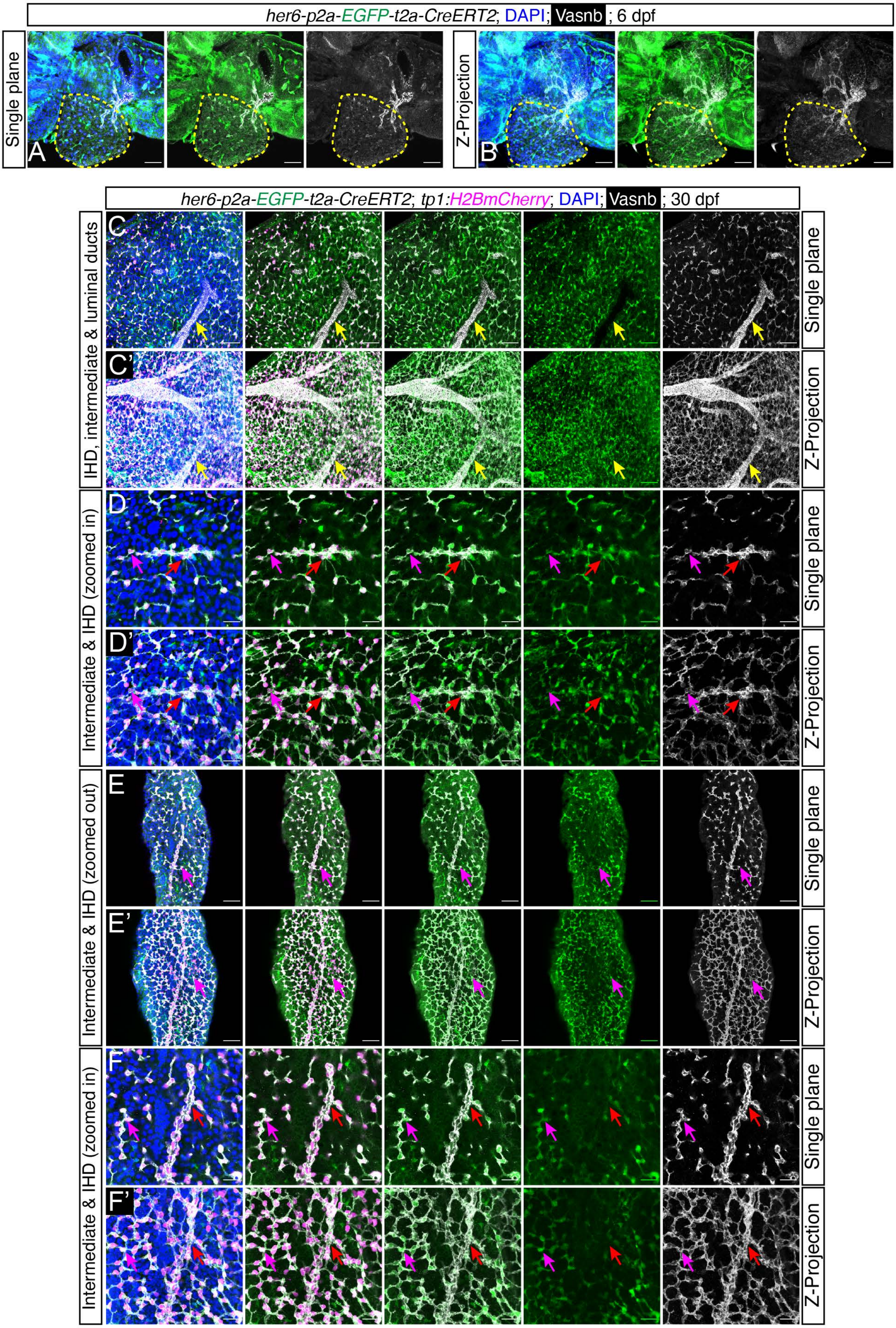
her6-p2a-EGFP-t2a-CreERT2; DAPI; Vasnb ; 6 dpf
Single plane
Z-Projection
A
B
her6-p2a-EGFP-t2a-CreERT2; tp1:H2BmCherry; DAPI; Vasnb ; 30 dpf
IHD, intermediate & luminal ducts
Intermediate & IHD (zoomed in)
Intermediate & IHD (zoomed out)
Intermediate & IHD (zoomed in)
C
C'
D
D'
E
E'
F
F'
Single plane
Z-Projection

**Supplementary Fig. 3. Immunostaining showing the *her6* expression pattern in hepatic biliary duct in larval and juvenile fish.** (A–B) Representative single-plane (A) or Z-projection (B) confocal images for the visualization of *her6* expression in zebrafish larvae using *TgKI(her6-p2A-EGFP-t2A-CreERT2)* homozygous knock-in reporter line. The yellow dashed line indicates the liver. (C–F) Representative single-plane (C,D,E,F) or Z-projection (C',D',E',F') confocal images for the visualization of *her6* expression pattern in juvenile zebrafish using *TgKI(her6-p2A-EGFP-t2A-CreERT2)* homozygous knock-in reporter line in combination with *Tg(tp1:H2BmCherry)*. Scale bars, 40 μm (A,B,D,F), 200 μm (C and E). The arrows denote the ducts with different morphology and spatial distribution, i.e., luminal duct (yellow) and intermediate duct (red), IHD (magenta).

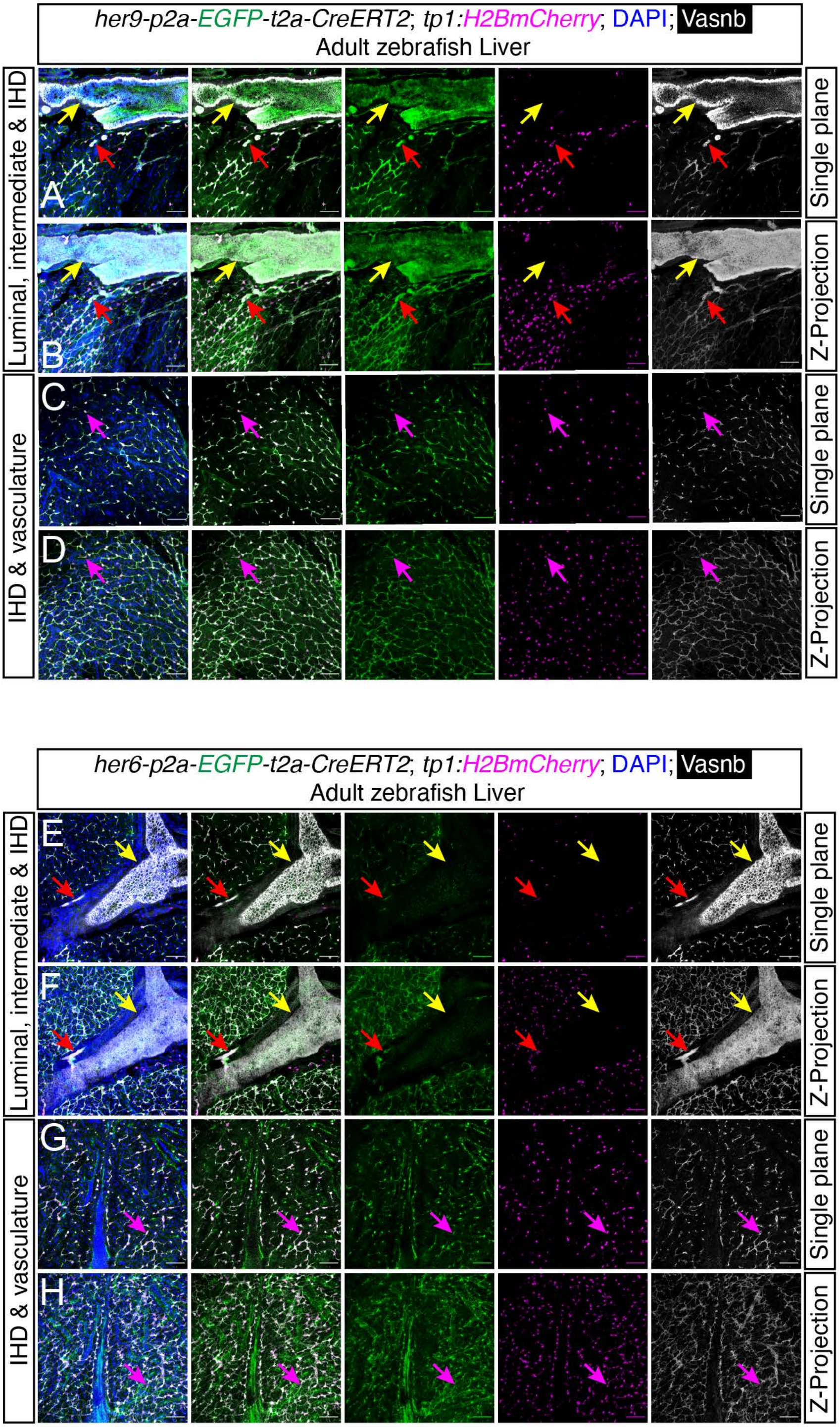
her9-p2a-EGFP-t2a-CreERT2; tp1:H2BmCherry; DAPI; Vasnb
Adult zebrafish Liver
Luminal, intermediate & IHD
IHD & vasculature
Single plane
Z-Projection
Single plane
Z-Projection
A
B
C
D
her6-p2a-EGFP-t2a-CreERT2; tp1:H2BmCherry; DAPI; Vasnb
Adult zebrafish Liver
Luminal, intermediate & IHD
IHD & vasculature
Single plane
Z-Projection
Single plane
Z-Projection
E
F
G
H

**Supplementary Fig. 4. *hes1* gene family expression in hepatic biliary system in adult zebrafish liver.** (A–D) Representative single-plane (A and C) or Z-projection (B and D) confocal images for the visualization of *her9* expression pattern in the hilum (A and B) and peripheral region (C and D) using adult *TgKI(her9-p2A-EGFP-t2A-CreERT2)* homozygous knock-in reporter line. (E–H) Representative single-plane (E and G) or Z-projection (F and H) confocal images for the visualization of *her6* expression pattern in the hilum (F and G) and peripheral region (H and I) using adult *TgKI(her6-p2A-EGFP-t2A-CreERT2)* homozygous knock-in reporter line. Scale bars, 200 μm. The arrows point to the ducts with different morphology and spatial distribution, i.e., luminal duct (yellow), intermediate duct (red), and IHD (magenta).

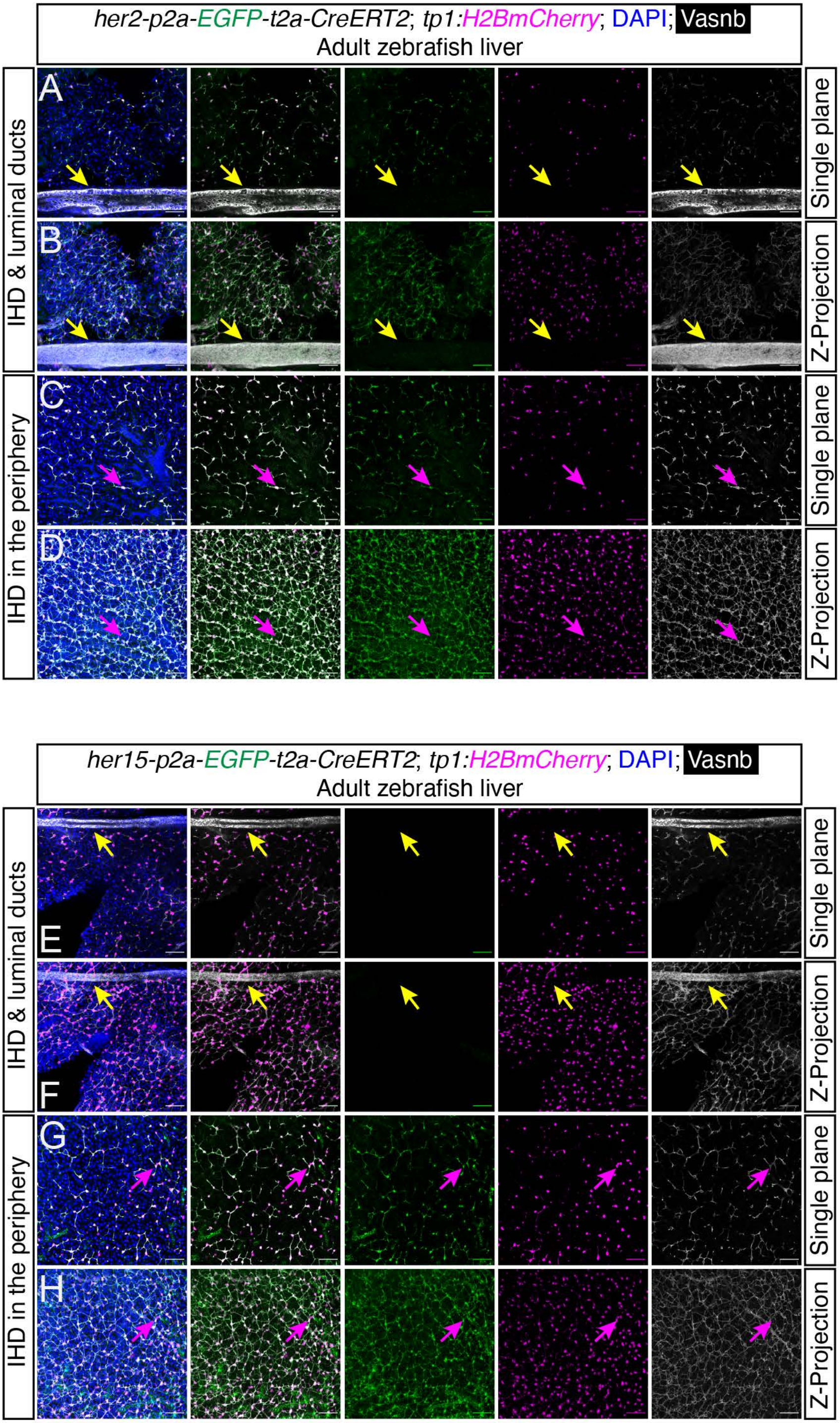

her2-p2a-EGFP-t2a-CreERT2; tp1:H2BmCherry; DAPI; Vasnb
Adult zebrafish liver
IHD & luminal ducts
IHD in the periphery
A
B
C
D
Single plane
Z-Projection
Single plane
Z-Projection
her15-p2a-EGFP-t2a-CreERT2; tp1:H2BmCherry; DAPI; Vasnb
Adult zebrafish liver
IHD & luminal ducts
IHD in the periphery
E
F
G
H
Single plane
Z-Projection
Single plane
Z-Projection

**Supplementary Fig. 5. *hes5* gene family expression in hepatic biliary system in adult zebrafish liver.** (A–D) Representative single-plane (A and C) or Z-projection (B and D) confocal images for the visualization of *her2* expression pattern in the hilum (A and B) and peripheral region (C and D) using adult *TgKI(her2-p2A-EGFP-t2A-CreERT2)* homozygous knock-in reporter line. (E–H) Representative single-plane (E and G) or Z-projection (F and H) confocal images for the visualization of *her15 (her15.1/her15.2)* expression pattern in the hilum (E and F) and peripheral region (G and H) using adult *TgKI(her15.1/15.2-p2A-EGFP-t2A-CreERT2)* homozygous knock-in reporter line. Scale bars, 200 μm. The arrows indicate the ducts with different morphology and spatial distribution, i.e., luminal duct (yellow) and IHD (magenta).

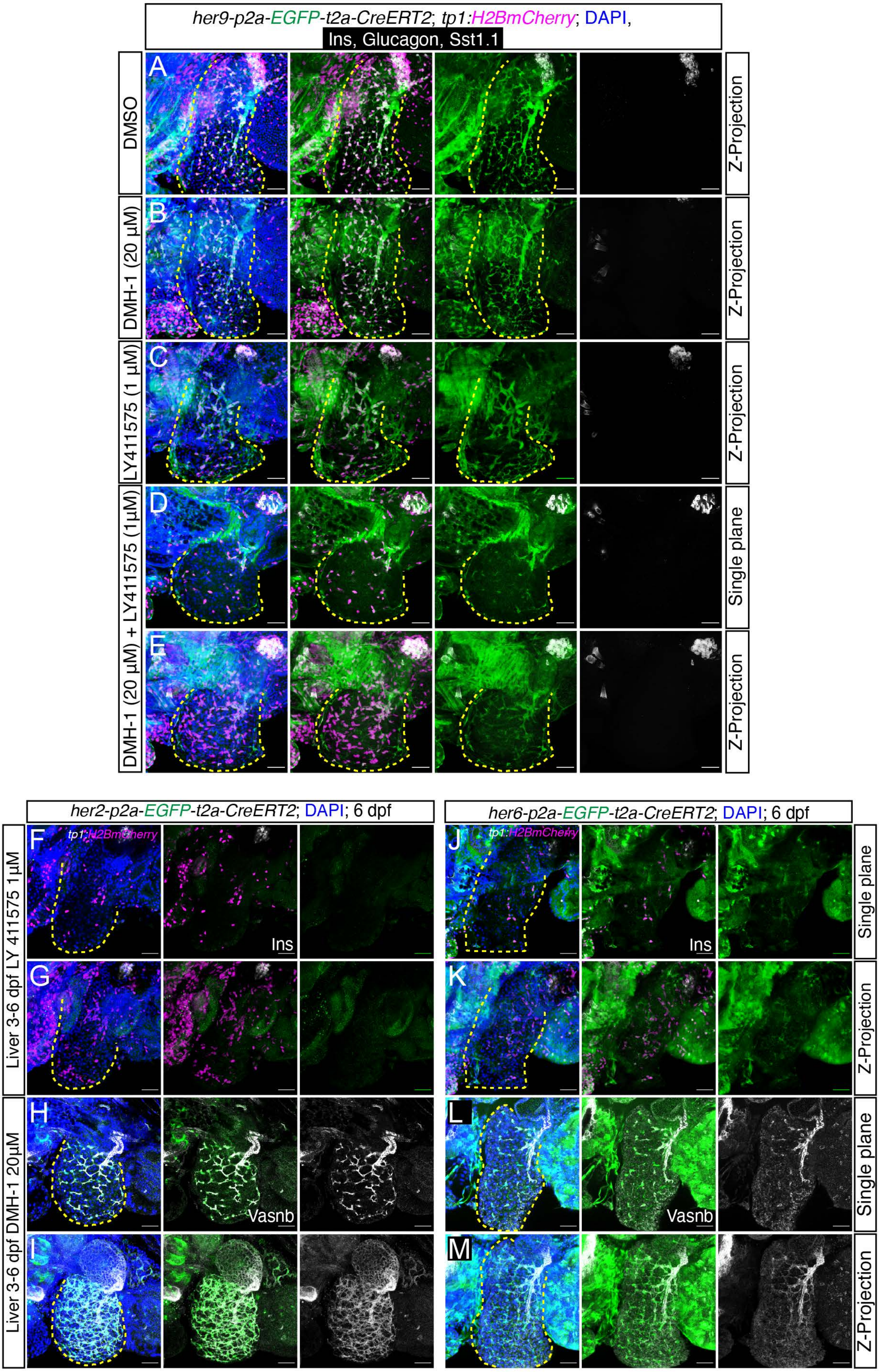

her9-p2a-EGFP-t2a-CreERT2; tp1:H2BmCherry; DAPI,
Ins, Glucagon, Sst1.1
DMSO
DMH-1 (20 μM)
LY411575 (1 μM)
DMH-1 (20 μM) + LY411575 (1μM)
Z-Projection
Single plane
her2-p2a-EGFP-t2a-CreERT2; DAPI; 6 dpf
her6-p2a-EGFP-t2a-CreERT2; DAPI; 6 dpf
tp1:H2BmCherry
Liver 3-6 dpf LY 411575 1μM
Liver 3-6 dpf DMH-1 20μM
Ins
Vasnb

**Supplementary Fig. 6. *her* gene family responsiveness to BMP and Notch signaling pathways.** (A–E) Representative Z-projection (A, B, C, E) or single-plane (D) confocal images following treatment with the control vehicle (DMSO), the BMP inhibitor (DMH-1), the Notch inhibitor (LY411575) or the combination of DMH-1 and LY411575 in *TgKI(her9-p2A-EGFP-t2A-CreERT2)* homozygous knock-in reporter line in the background of *Tg(tp1:H2BmCherry)*. (F-I) Representative single-plane (F and H) or Z-projection (G and I) confocal images following treatment with either the Notch inhibitor (LY411575) or the BMP inhibitor (DMH-1) in *TgKI(her2-p2A-EGFP-t2A-CreERT2)* homozygous knock-in reporter line. (J-M) Representative single-plane (J and L) or Z-projection (K and M) confocal images following treatment with either the Notch inhibitor (LY411575) or the BMP inhibitor (DMH-1) in *TgKI(her6-p2A-EGFP-t2A-CreERT2)* homozygous knock-in reporter line. Untreated *TgKI(her2-p2A-EGFP-t2A-CreERT2)* and *TgKI(her6-p2A-EGFP-t2A-CreERT2)* larva can be viewed in Figure 2B and S3A-B, respectively. Scale bars, 100 µm. The yellow dashed lines indicate the liver.

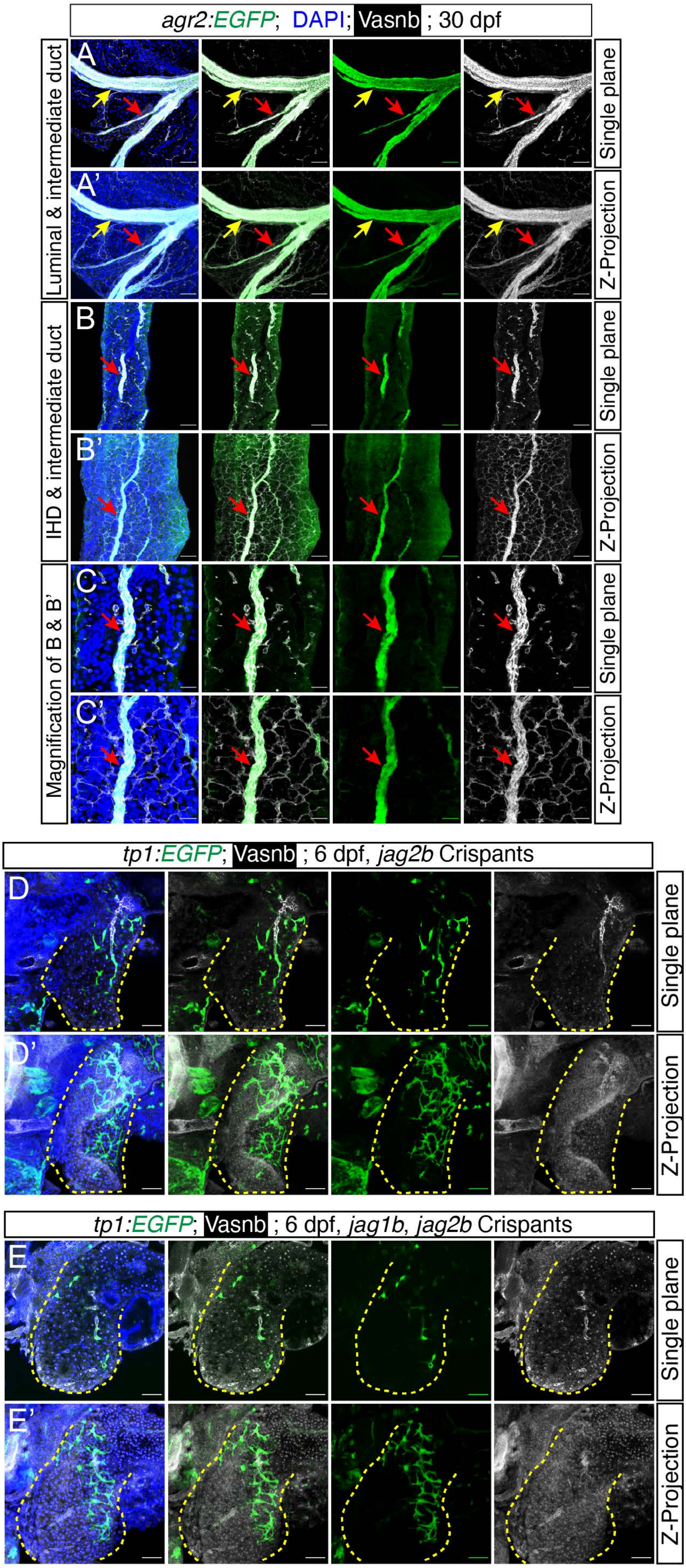

agr2:EGFP; DAPI; Vasnb ; 30 dpf
Luminal & intermediate duct
IHD & intermediate duct
Magnification of B & B'
A
A'
B
B'
C
C'
Single plane
Z-Projection
tp1:EGFP; Vasnb ; 6 dpf, jag2b Crispants
D
D'
tp1:EGFP; Vasnb ; 6 dpf, jag1b, jag2b Crispants
E
E'

**Supplementary Fig. 7. Duct heterogeneity and ductal cell plasticity.** (A-C) Immunostaining in *Tg(agr2:EGFP)* juveniles showing the ducts in the hepatic biliary system with representative single-plane (A, B, C) or Z-projection (A', B', C') confocal images. (D and E) Single-plane (D, E) or Z-projection (D', E') confocal images showing the labeling pattern in *Tg(tp1:EGFP)* with *jag2b* single or *jag1b*/*jag2b* double CRISPR-Cas9 mediated mutagenesis. Scale bars, 200 μm (A and B); 40 μm (C - E). The arrows point to the ducts with different morphology and spatial distribution, i.e., luminal duct (yellow) and intermediate duct (red). The yellow dashed lines outline the liver.

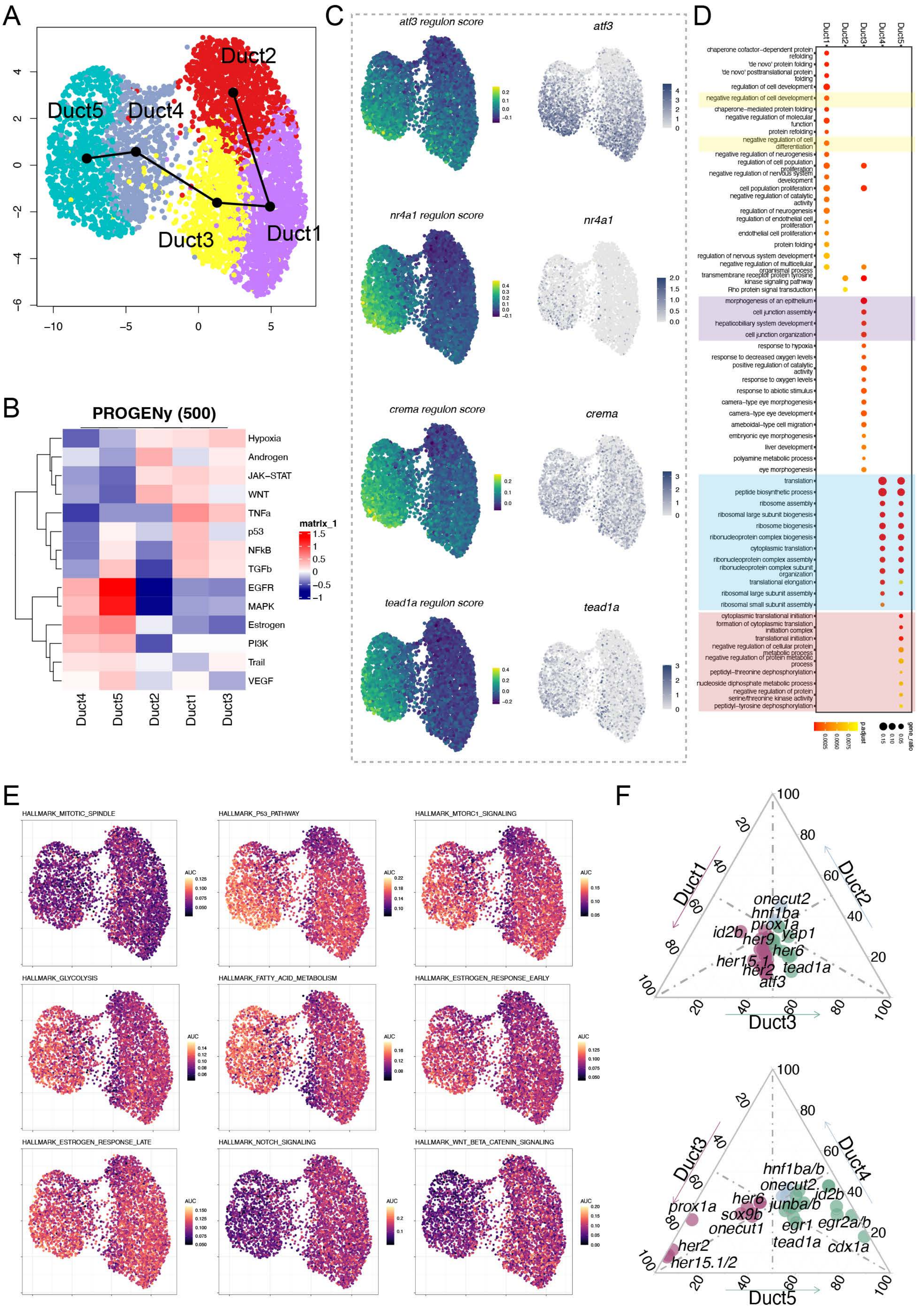
A
Duct2
Duct5
Duct4
Duct3
Duct1
B
PROGENy (500)
Hypoxia
Androgen
JAK–STAT
WNT
TNFa
p53
NFkB
TGFb
EGFR
MAPK
Estrogen
PI3K
Trail
VEGF
matrix_1
Duct4
Duct5
Duct2
Duct1
Duct3
C
atf3 regulon score
atf3
nr4a1 regulon score
nr4a1
crema regulon score
crema
tead1a regulon score
tead1a
D
Duct1
Duct2
Duct3
Duct4
Duct5
chaperone cofactor–dependent protein refolding
'de novo' protein folding
'de novo' posttranslational protein folding
regulation of cell development
negative regulation of cell development
chaperone–mediated protein folding
negative regulation of molecular function
protein refolding
negative regulation of cell differentiation
negative regulation of neurogenesis
regulation of cell population proliferation
negative regulation of nervous system development
cell population proliferation
negative regulation of catalytic activity
regulation of neurogenesis
regulation of endothelial cell proliferation
endothelial cell proliferation
protein folding
regulation of nervous system development
negative regulation of multicellular organismal process
transmembrane receptor protein tyrosine kinase signaling pathway
Rho protein signal transduction
morphogenesis of an epithelium
cell junction assembly
hepaticobiliary system development
cell junction organization
response to hypoxia
response to decreased oxygen levels
positive regulation of catalytic activity
response to oxygen levels
response to abiotic stimulus
camera–type eye morphogenesis
camera–type eye development
ameboidal–type cell migration
embryonic eye morphogenesis
liver development
polyamine metabolic process
eye morphogenesis
translation
peptide biosynthetic process
ribosome assembly
ribosomal large subunit biogenesis
ribosome biogenesis
ribonucleoprotein complex biogenesis
cytoplasmic translation
ribonucleoprotein complex assembly
ribonucleoprotein complex subunit organization
translational elongation
ribosomal large subunit assembly
ribosomal small subunit assembly
cytoplasmic translational initiation
formation of cytoplasmic translation initiation complex
translational initiation
negative regulation of cellular protein metabolic process
negative regulation of protein metabolic process
peptidyl–threonine dephosphorylation
nucleoside diphosphate metabolic process
negative regulation of protein serine/threonine kinase activity
peptidyl–tyrosine dephosphorylation
p.adjust
gene_ratio
E
HALLMARK_MITOTIC_SPINDLE
HALLMARK_P53_PATHWAY
HALLMARK_MTORC1_SIGNALING
HALLMARK_GLYCOLYSIS
HALLMARK_FATTY_ACID_METABOLISM
HALLMARK_ESTROGEN_RESPONSE_EARLY
HALLMARK_ESTROGEN_RESPONSE_LATE
HALLMARK_NOTCH_SIGNALING
HALLMARK_WNT_BETA_CATENIN_SIGNALING
AUC
F
Duct1
Duct2
Duct3
onecut2
hnf1ba
prox1a
id2b
yap1
her9
her6
her15.1
her2
tead1a
atf3
Duct3
Duct4
Duct5
hnf1ba/b
onecut2
her6
id2b
junba/b
sox9b
prox1a
onecut1
egr1
egr2a/b
tead1a
cdx1a
her2
her15.1/2

**Supplementary Fig. 8. *In silico* analyses of hepatic ductal cells in juvenile fish liver.** (A) Slingshot pseudotime trajectory showing the cell transition flow among different liver duct subclusters, starting with Duct2 as the root. (B) PROGENy analysis demonstrating the pathway activity in different cell clusters. (C) The UMAP for the developing ductal cells based on the RASs (left) next to the expression level of the transcription factors (right). (D) Functional GO enrichment indicating the significant GO terms for each ductal cell cluster. (E) UMAP plot showing the statistically significant gene activity scores calculated by AUCell for each cell. (F) Ternary plot shows the expression of DNA binding genes in pairs of Duct1/2/3 and pairs of Duct3/4/5. RAS: regulon activity score. GO: Gene Ontology.

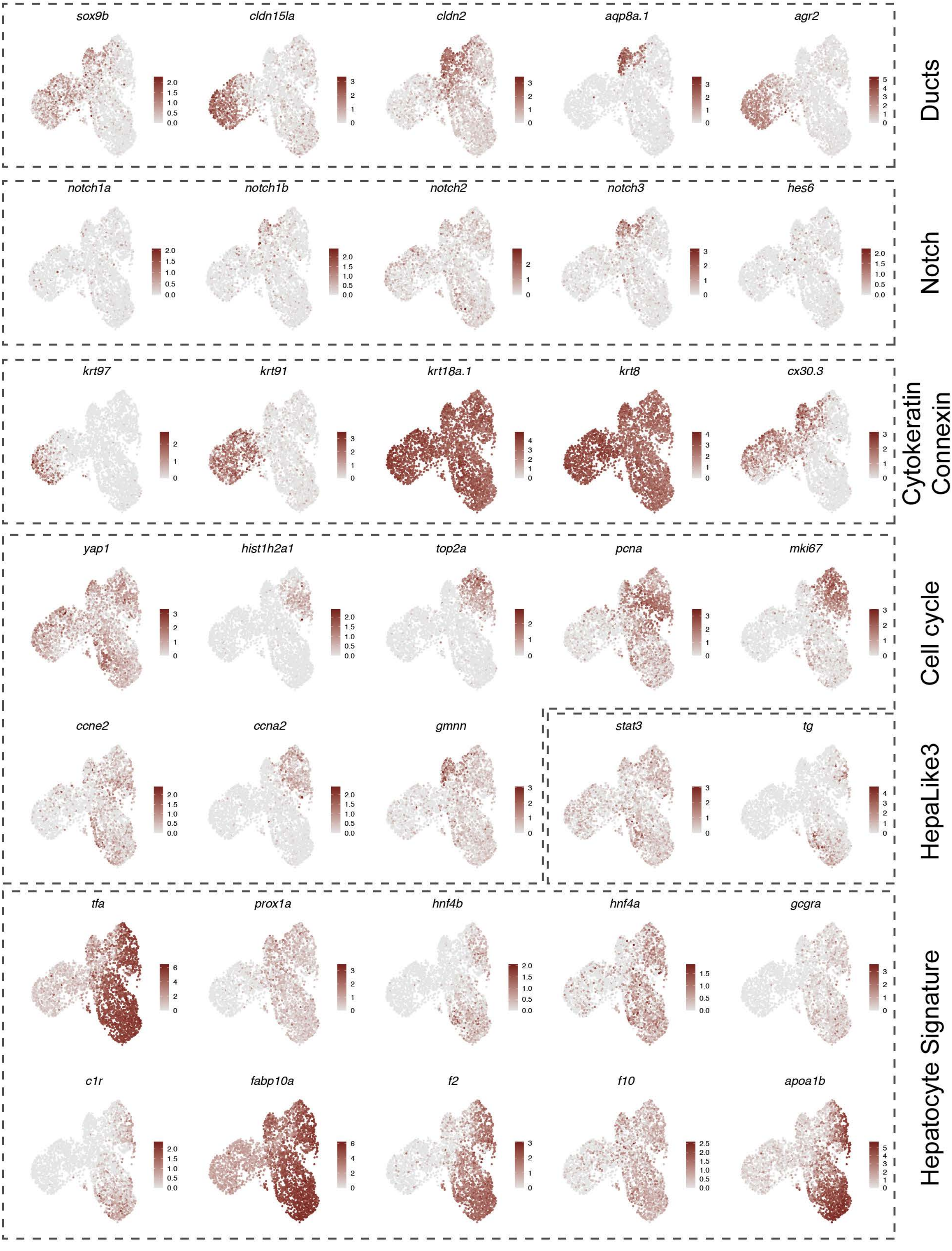
sox9b
cldn15la
cldn2
aqp8a.1
agr2
Ducts
notch1a
notch1b
notch2
notch3
hes6
Notch
krt97
krt91
krt18a.1
krt8
cx30.3
Cytokeratin
Connexin
yap1
hist1h2a1
top2a
pcna
mki67
Cell cycle
ccne2
ccna2
gmnn
stat3
tg
HepaLike3
tfa
prox1a
hnf4b
hnf4a
gcgra
c1r
fabp10a
f2
f10
apoa1b
Hepatocyte Signature

**Supplementary Fig. 9. Single-cell transcriptomics highlight distinct molecular signatures in various cell clusters during liver regeneration.** UMAP plots of various marker genes, colored by the normalized gene expression levels.

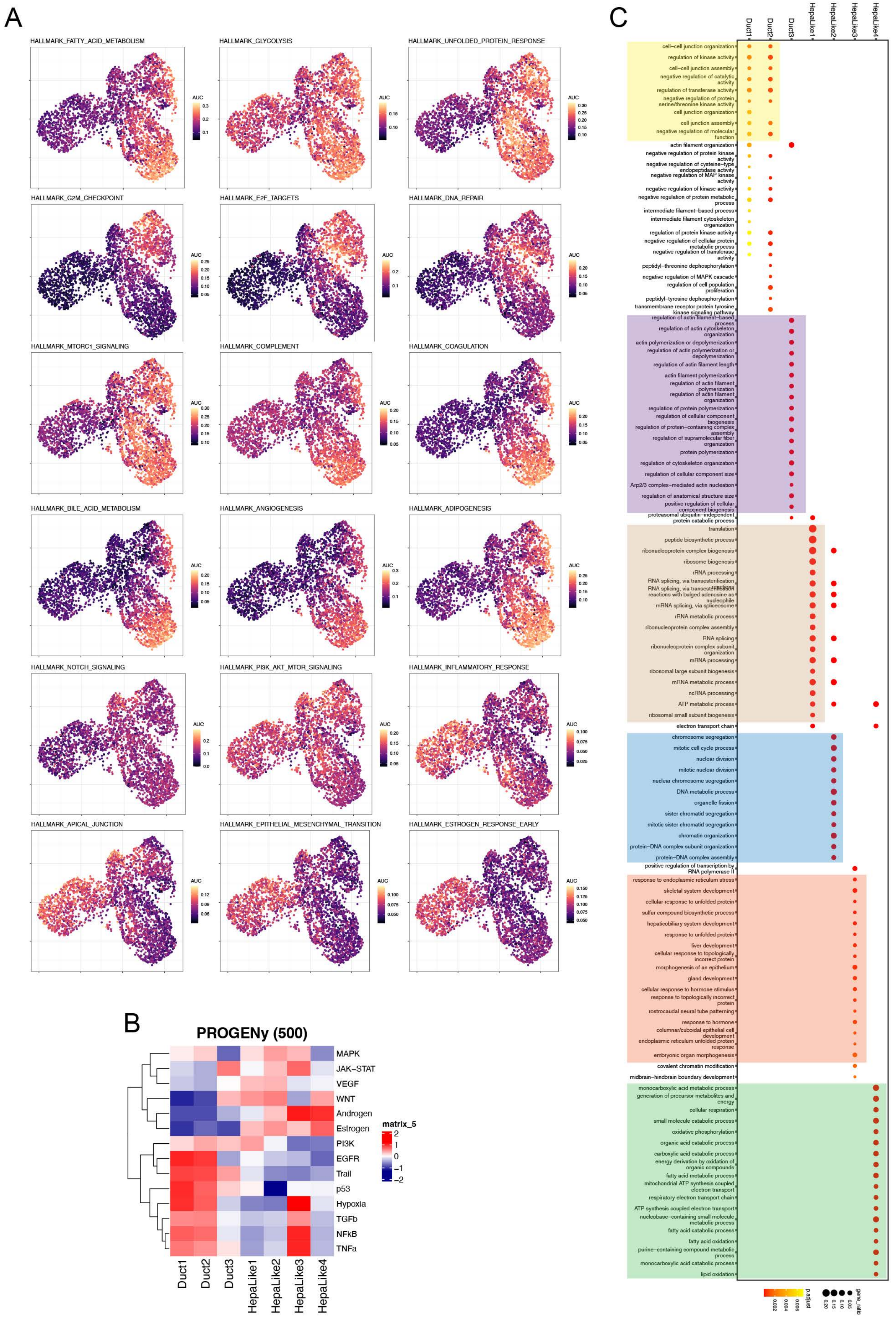

A
HALLMARK_FATTY_ACID_METABOLISM
HALLMARK_GLYCOLYSIS
HALLMARK_UNFOLDED_PROTEIN_RESPONSE
HALLMARK_G2M_CHECKPOINT
HALLMARK_E2F_TARGETS
HALLMARK_DNA_REPAIR
HALLMARK_MTORC1_SIGNALING
HALLMARK_COMPLEMENT
HALLMARK_COAGULATION
HALLMARK_BILE_ACID_METABOLISM
HALLMARK_ANGIOGENESIS
HALLMARK_ADIPOGENESIS
HALLMARK_NOTCH_SIGNALING
HALLMARK_PI3K_AKT_MTOR_SIGNALING
HALLMARK_INFLAMMATORY_RESPONSE
HALLMARK_APICAL_JUNCTION
HALLMARK_EPITHELIAL_MESENCHYMAL_TRANSITION
HALLMARK_ESTROGEN_RESPONSE_EARLY
AUC
B
PROGENy (500)
MAPK
JAK–STAT
VEGF
WNT
Androgen
Estrogen
PI3K
EGFR
Trail
p53
Hypoxia
TGFb
NFkB
TNFa
matrix_5
2
1
0
−1
−2
Duct1
Duct2
Duct3
HepaLike1
HepaLike2
HepaLike3
HepaLike4
C
Duct1
Duct2
Duct3
HepaLike1
HepaLike2
HepaLike3
HepaLike4
cell–cell junction organization
regulation of kinase activity
cell–cell junction assembly
negative regulation of catalytic activity
regulation of transferase activity
negative regulation of protein serine/threonine kinase activity
cell junction organization
cell junction assembly
negative regulation of molecular function
actin filament organization
negative regulation of protein kinase activity
negative regulation of cysteine–type endopeptidase activity
negative regulation of MAP kinase activity
negative regulation of kinase activity
negative regulation of protein metabolic process
intermediate filament–based process
intermediate filament cytoskeleton organization
regulation of protein kinase activity
negative regulation of cellular protein metabolic process
negative regulation of transferase activity
peptidyl–threonine dephosphorylation
negative regulation of MAPK cascade
regulation of cell population proliferation
peptidyl–tyrosine dephosphorylation
transmembrane receptor protein tyrosine kinase signaling pathway
regulation of actin filament–based process
regulation of actin cytoskeleton organization
actin polymerization or depolymerization
regulation of actin polymerization or depolymerization
regulation of actin filament length
actin filament polymerization
regulation of actin filament polymerization
regulation of actin filament organization
regulation of protein polymerization
regulation of cellular component biogenesis
regulation of protein–containing complex assembly
regulation of supramolecular fiber organization
protein polymerization
regulation of cytoskeleton organization
regulation of cellular component size
Arp2/3 complex–mediated actin nucleation
regulation of anatomical structure size
positive regulation of cellular component biogenesis
proteasomal ubiquitin–independent protein catabolic process
translation
peptide biosynthetic process
ribonucleoprotein complex biogenesis
ribosome biogenesis
rRNA processing
RNA splicing, via transesterification reactions
RNA splicing, via transesterification reactions with bulged adenosine as nucleophile
mRNA splicing, via spliceosome
rRNA metabolic process
ribonucleoprotein complex assembly
RNA splicing
ribonucleoprotein complex subunit organization
mRNA processing
ribosomal large subunit biogenesis
mRNA metabolic process
ncRNA processing
ATP metabolic process
ribosomal small subunit biogenesis
electron transport chain
chromosome segregation
mitotic cell cycle process
nuclear division
mitotic nuclear division
nuclear chromosome segregation
DNA metabolic process
organelle fission
sister chromatid segregation
mitotic sister chromatid segregation
chromatin organization
protein–DNA complex subunit organization
protein–DNA complex assembly
positive regulation of transcription by RNA polymerase II
response to endoplasmic reticulum stress
skeletal system development
cellular response to unfolded protein
sulfur compound biosynthetic process
hepaticobiliary system development
response to unfolded protein
liver development
cellular response to topologically incorrect protein
morphogenesis of an epithelium
gland development
cellular response to hormone stimulus
response to topologically incorrect protein
rostrocaudal neural tube patterning
response to hormone
columnar/cuboidal epithelial cell development
endoplasmic reticulum unfolded protein response
embryonic organ morphogenesis
covalent chromatin modification
midbrain–hindbrain boundary development
monocarboxylic acid metabolic process
generation of precursor metabolites and energy
cellular respiration
small molecule catabolic process
oxidative phosphorylation
organic acid catabolic process
carboxylic acid catabolic process
energy derivation by oxidation of organic compounds
fatty acid metabolic process
mitochondrial ATP synthesis coupled electron transport
respiratory electron transport chain
ATP synthesis coupled electron transport
nucleobase–containing small molecule metabolic process
fatty acid catabolic process
fatty acid oxidation
purine–containing compound metabolic process
monocarboxylic acid catabolic process
lipid oxidation
p.adjust
gene_ratio

**Supplementary Fig. 10. Functional enrichment of cell clusters in liver regeneration dataset.** (A) UMAP plot showing the statistically significant gene activity scores calculated by AUCell for each cell. (B) PROGENy analysis demonstrating the pathway activity in different cell clusters. (C) Functional GO enrichment indicating the significant GO terms for each cell cluster. AUC, area under the curve. GO: Gene Ontology.

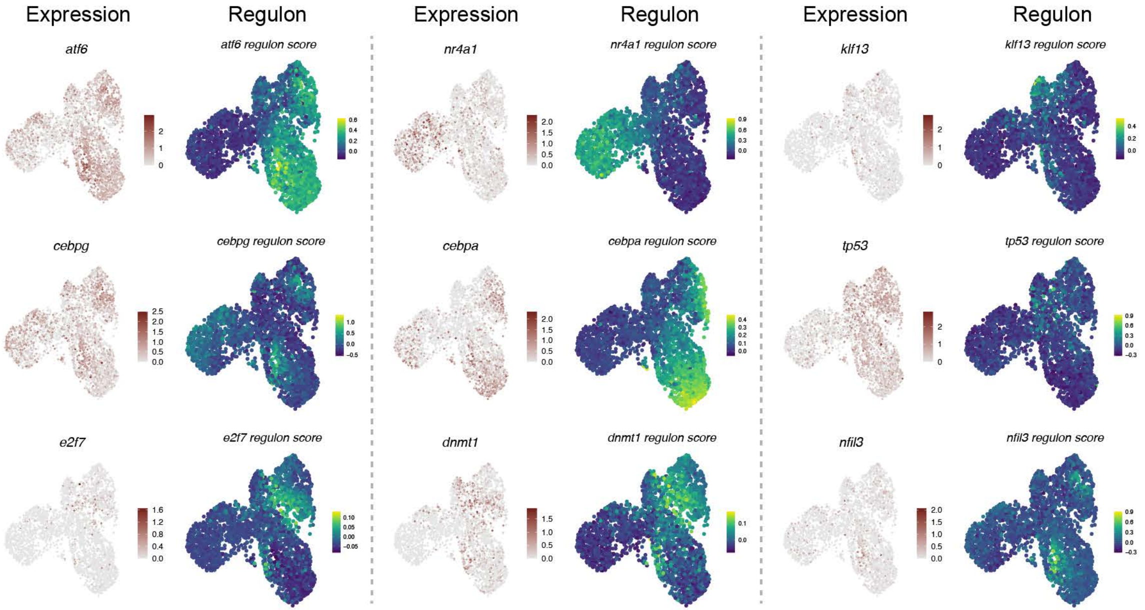
Expression
Regulon
Expression
Regulon
Expression
Regulon
atf6
atf6 regulon score
nr4a1
nr4a1 regulon score
klf13
klf13 regulon score
cebpg
cebpg regulon score
cebpa
cebpa regulon score
tp53
tp53 regulon score
e2f7
e2f7 regulon score
dnmt1
dnmt1 regulon score
nfil3
nfil3 regulon score

**Supplementary Fig. 11. *In silico* SCENIC regulon analyses during liver regeneration.** The UMAP for regenerative ductal and HepaLike cells based on the RASs (right), next to the expression level of the transcription factors (left). RAS: regulon activity scores.
